# Quantitative high-throughput tests of ubiquitous RNA secondary structure prediction algorithms via RNA/protein binding

**DOI:** 10.1101/571588

**Authors:** Winston R. Becker, Inga Jarmoskaite, Kalli Kappel, Pavanapuresan P. Vaidyanathan, Sarah K. Denny, Rhiju Das, William J. Greenleaf, Daniel Herschlag

## Abstract

Nearest-neighbor (NN) rules provide a simple and powerful quantitative framework for RNA structure prediction that is strongly supported for canonical Watson-Crick duplexes from a plethora of thermodynamic measurements. Predictions of RNA secondary structure based on nearest-neighbor (NN) rules are routinely used to understand biological function and to engineer and control new functions in biotechnology. However, NN applications to RNA structural features such as internal and terminal loops rely on approximations and assumptions, with sparse experimental coverage of the vast number of possible sequence and structural features. To test to what extent NN rules accurately predict thermodynamic stabilities across RNAs with non-WC features, we tested their predictions using a quantitative high-throughput assay platform, RNA-MaP. Using a thermodynamic assay with coupled protein binding, we carried out equilibrium measurements for over 1000 RNAs with a range of predicted secondary structure stabilities. Our results revealed substantial scatter and systematic deviations between NN predictions and observed stabilities. Solution salt effects and incorrect or omitted loop parameters contribute to these observed deviations. Our results demonstrate the need to independently and quantitatively test NN computational algorithms to identify their capabilities and limitations. RNA-MaP and related approaches can be used to test computational predictions and can be adapted to obtain experimental data to improve RNA secondary structure and other prediction algorithms.

**Significance statement:** RNA secondary structure prediction algorithms are routinely used to understand, predict and design functional RNA structures in biology and biotechnology. Given the vast number of RNA sequence and structural features, these predictions rely on a series of approximations, and independent tests are needed to quantitatively evaluate the accuracy of predicted RNA structural stabilities. Here we measure the stabilities of over 1000 RNA constructs by using a coupled protein binding assay. Our results reveal substantial deviations from the RNA stabilities predicted by popular algorithms, and identify factors contributing to the observed deviations. We demonstrate the importance of quantitative, experimental tests of computational RNA structure predictions and present an approach that can be used to routinely test and improve the prediction accuracy.

## Introduction

Structure is foundational to RNA function, from post-transcriptional regulation by RNA aptamers and small RNAs to catalysis by intricate RNA enzymes. RNA secondary structure constitutes the ‘backbone’ of structured RNAs, contributing the bulk of structural stability. The ability of mRNAs and other RNAs to associate with proteins, other RNAs, and small molecules is facilitated or repressed by the formation of RNA structure. Correspondingly, predictive models of RNA secondary structure energetics are indispensable to understand RNA biology, in therapeutic development, and in design. The widespread use of secondary structure prediction algorithms is reflected in >10,000 citations (Table S1), with applications spanning the discovery of functional RNA structures in biology, mechanistic dissection of RNA structure-function relationships, the design of small RNAs for RNAi and CRISPR, and RNA nanotechnology (e.g.,(1-9)).

Nearest-neighbor (NN) algorithms provide a powerful and simple framework for RNA structure predictions. The NN model originates from the early 1970s, when Uhlenbeck, Tinoco, Crothers and others demonstrated that the stability of a structured RNA is well described by an additive free energy model based on the stabilities of neighboring RNA base pairs and intervening loops (10). Hundreds of thermodynamic parameters have since been determined for the different base-pair neighbors and types and sequences of RNA loops, with the most recent parameter set (‘Turner 2004’) comprised of 294 parameters derived from 802 thermodynamic measurements (11, 12). However, the extent of experimental coverage and quality of predictions varies widely between structural features. For example, all 10 possible canonical base-pair neighbors have been measured in multiple contexts, and consequently RNA duplex stabilities can be predicted with high accuracy (13). In contrast, only a small fraction (<10%) of the 1024^1^ possible tetraloops have been measured (11, 14), with several displaying exceptionally low or high stabilities that span a range of 5 kcal/mol at 37 °C. The coverage becomes even lower for features such as asymmetric internal loops, despite their common occurrence in biological RNAs: e.g., fewer than 0.1% of possible 3×5 internal loops sequences have been measured (7 of >10^4^)(15).

In an effort to expand the NN tests beyond the limited thermodynamic measurements, NN prediction algorithms are now routinely tested, and in some cases parametrized, on the basis of RNA structure databases that comprise thousands of experimental RNA structures (16-18). In this approach, the fraction of base pairs that do or do not occur in the structure are evaluated and optimized by adjusting the algorithm. However, a correctly predicted structure may have a stability of –1 kcal/mol or –100 kcal/mol, while a structure scored as ‘incorrect’ may be only marginally (<1 kcal/mol) less stable than a competing structure, and it is unclear how to fully account for such marginal stability in this type of analysis (18); thus, evaluation of structures as either correct or incorrect is likely not suitable for quantitatively assessing NN rules. Furthermore, the structures in the RNA database may be confounded by tertiary structure and protein binding, which occur abundantly in the large RNAs and RNA/protein complexes that dominate the database (e.g., ribosomal RNAs, self-splicing introns, tRNAs, RNAse P). Thus, while attempts to incorporate structural data to make up for the lack of experimentally defined parameters are important, they are inherently limited, and quantitative thermodynamic measurements of well-defined RNA secondary structure constructs are needed to parameterize and test these algorithms.

Advances in high-throughput biophysics now make it possible to systematically probe energetics for large numbers of RNAs (19-21). The RNA-MaP platform allows direct, parallel equilibrium and kinetic measurements of binding of thousands of RNAs with their RNA or protein interaction partners. Here we used RNA-MaP to determine the stabilities of a library of over 1000 RNA hairpin constructs, by quantifying the extent to which structure formation inhibits binding of the well-characterized single-stranded RNA binding proteins PUM1 and PUM2 (22-27). Our results reveal substantial and systematic deviations between structure stabilities predicted by current algorithms and measured stabilities. These observations underscore the value of blind tests and the need for additional parametrization of these secondary structure algorithms.

## Results and Discussion

### Coupled equilibrium binding to quantify RNA structural stability

The coupled equilibrium binding approach is shown schematically in Fig. 1A. An RBP, in our case PUM1/2, binds its specific RNA site in an unstructured, base-pair free state. RNA secondary structure involving residues in the RBP site decreases the site accessibility, thereby lowering the observed affinity to an extent proportional to the stability of RNA structure (Fig. 1A). By measuring equilibrium binding of the RBP to a library of structured RNA sequences and by comparing the resulting affinities with PUM2 affinity for its unstructured consensus sequence, the stability of RNA structure blocking the PUM2 site is determined (Fig. 1A).

**Fig. 1.**
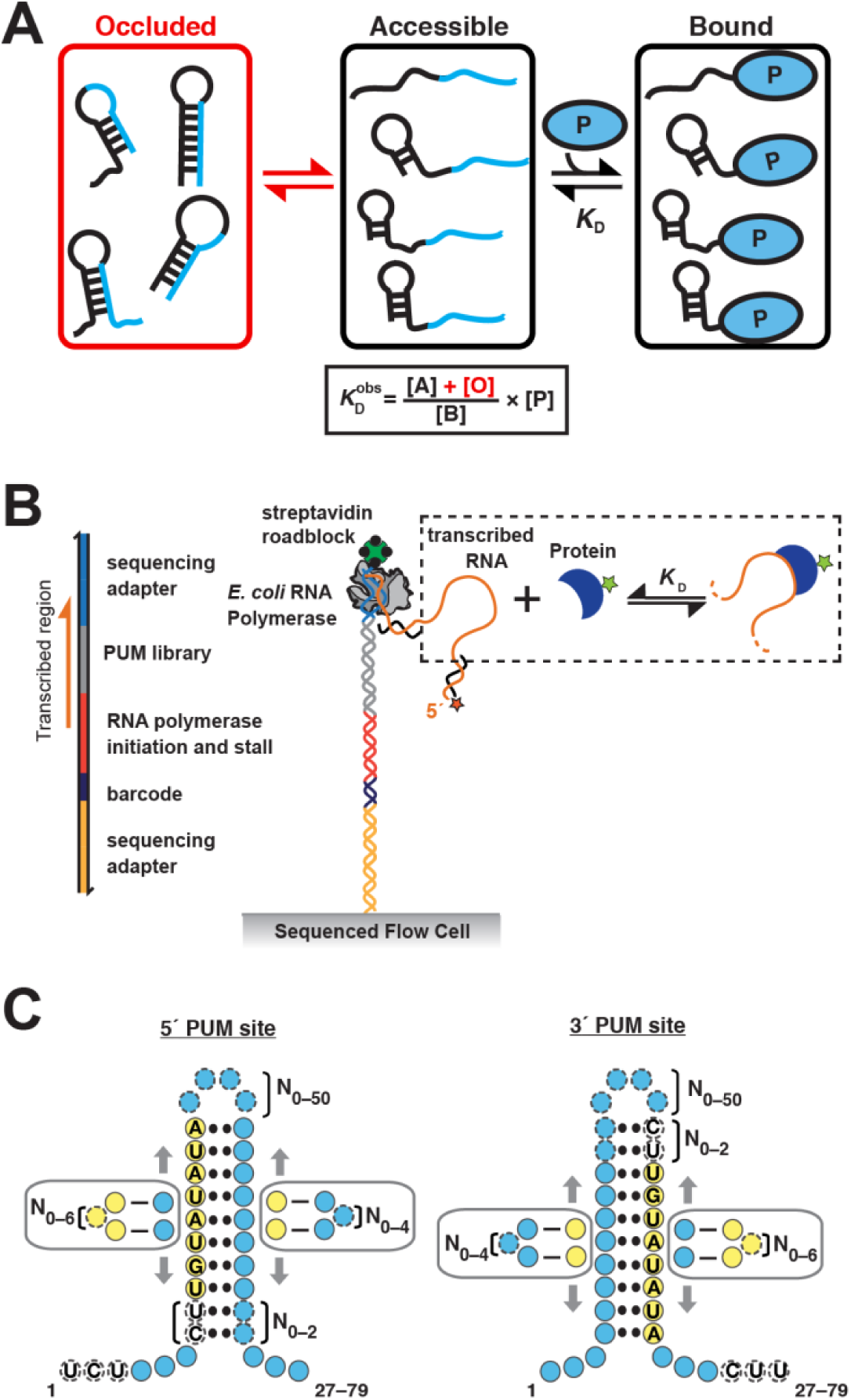
Testing RNA structure stability via PUM2 binding. (A) RNA structure renders the binding site of a single-stranded RNA binding protein inaccessible thereby increasing the observed dissociation constant 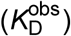 to an extent that reflects the stability of RNA structure (reproduced from (27)). (B) RNA-MaP is used to measure protein binding to RNA in high throughput (19). *Left*: Schematic representation of RNA-MaP constructs used in this study. *Right*: Schematic depiction of an RNA-MaP experiment. The “PUM library” RNA, with short constant flanking sequences, was transcribed on a sequenced MiSeq flow cell, and RNA transcripts were immobilized at the end of their DNA template by stalling the RNA polymerase with a terminal streptavidin block. RNA was visualized by hybridization of its constant 5’-terminal sequence to a fluorescently labeled DNA oligonucleotide (red star). Fluorescently-labeled protein was then flowed at a series of increasing concentrations, and following equilibration, the amount of protein co-localized with each RNA cluster at each concentration was determined in a different fluorescence channel (adapted from (27)). (C) Summary of the oligonucleotide library designs. Each base pair is present in a subset of constructs, as indicated with two dots. Circles indicate individual nucleotides, using the following color scheme: *yellow*, PUM1/2 binding site; *blue*, positions with variable sequence; *white*, positions with invariable sequence. Dashed outlines indicate nucleotides that are present only in a subset of constructs. The boxed nucleotides represent bulges, formed within or opposite from the PUM1/2 site, and the vertical arrows indicate varying bulge positions along the stem. The total length of the oligonucleotides ranged from 27 to 79 nt. See Fig. S1 for additional details and for constructs containing mutated PUM1/2 sites.

The coupled equilibrium binding approach provides an alternative to the traditionally employed UV optical melting measurements. In UV melting, the absorbance of an RNA oligonucleotide (or, in case of bimolecular interactions, two oligonucleotides) is measured across a range of temperatures, and the stability is determined by fitting the resulting curve to a two-state model, based on the absorbance differences between the unfolded and folded RNA. While powerful, optical melting measurements have limitations. Their interpretation typically relies on two-state and other simplifying assumptions (28). Further, RNA constructs that provide the most accurate thermodynamic measurements have melting temperatures above physiological, thus requiring significant extrapolation for biological application. In contrast, by quantifying the overall accessibility of RNA, the coupled equilibrium approach reports on the stability of the ensemble of RNA structures without assuming two-state behavior or making other assumptions about the composition or homogeneity of the folded state. While in principle indirect, the coupled equilibrium binding approach provides a quantitative readout of RNA structural stability at temperatures and conditions of interest. As a result, while we can envision multiple schema for adapting melting measurements for high-throughput studies, the approach used herein has the advantage of providing a fully independent test and avoiding the assumptions inherent to UV melting measurements.

The model underlying the coupled binding approach is derived from statistical thermodynamics and is applicable to any RNA-binding protein (RBP) that meets the following conditions. First, the model assumes no RBP binding to structurally occluded binding sites. Thus, the RBP must bind single-stranded RNA with affinity at least several orders of magnitude greater than any specific or non-specific binding to structured RNA. Second, the model assumes that the RBP binds accessible sites with constant affinity regardless of any RNA structure outside the binding site (Alternatively, RNAs that lack predicted persistent structure in the bound state can be used in analyses.) Third, the RBP must have known sequence specificity, to pinpoint the RNA residues whose accessibility determines affinity. We test and validate each of these premises below.

### Design of an RNA library to test NN rules

To provide an initial independent quantitative test of NN predictions, we designed a library of RNA hairpins containing single PUM1/2 binding sites (Fig. 1C, Fig. S1). We individually and in combination varied the number of base-pairs, the length and sequence of the hairpin loop, and the sequence, number and position of internal loops (Fig. S1). The position and sequence of the PUM1/2 sites were also varied. This design enabled testing NN predictions across sequence and structural features that might be better or worse described by NN algorithms. The ensemble stabilities predicted for the oligonucleotide using Vienna RNAfold (29) spanned a range of 11.1 and 8.3 kcal/mol at 25 and 37 °C, respectively. The library also included sequences with no predicted structure that served as references to quantify the effects of structure.

### Validation of a high-throughput assay for RNA structural stability

Recently-developed technologies like RNA-MaP allow the folding of thousands of different RNA sequences to be measured in parallel (19-21). In RNA-MaP, a library of RNAs is expressed and immobilized within sequence-identified clusters on an Illumina MiSeq chip, and binding of fluorescently-labeled interaction partners is directly monitored across a range of concentrations to determine binding affinity (Fig. 1B). To quantify the stabilities of RNA structures in our library, we used the RNA-MaP platform to measure PUM2 equilibrium binding to the oligonucleotide library. The data were fit to a single-site binding model with an additional binding term for non-specific PUM2 association with a PUM2/RNA complex (27). Representative binding curves for varying RNA stabilities are shown in Fig. S2. The binding affinities were highly reproducible (R^2^ = 0.96), and the median uncertainty was 0.21 kcal/mol, enabling high-precision tests of structure predictions (Fig. S3). The high-confidence data discussed below were based on variants with at least five measurements and a 95% confidence interval of measured affinity ≤1 kcal/mol. (See SI Materials and Methods and Ref. (27) for data fitting and quality assessment.)

We first established that PUM2 binding met the criteria for the simple thermodynamic model described in Fig. 1A. To assess the requirement for discrimination between single-stranded and structured binding sites, we compared PUM2 binding to consensus sites predicted to have no structure and sites with the most stable predicted structures, as determined by RNAfold (29). As expected, given the extensive structural evidence for recognition of RNA bases by PUM2 and its homologs, PUM2 showed negligible binding to sites occluded by stable structures 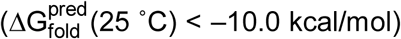, with affinities at least 10^4^-fold (5.4 kcal/mol) weaker than RNAs with an unstructured consensus sequence (Fig. S2).

The second condition is that binding affinity to accessible (i.e., non-base paired) sites is constant regardless of structure surrounding the accessible PUM2 binding site (Fig. 1A)—i.e., certain structures persist upon protein binding without affecting binding affinity. If nearby structures did affect protein binding due to steric or topological effects, we would most simply expect greater structure effects than predicted based on the accessibility of the PUM2 site. This behavior was observed only for a few variants with stable predicted structures in close vicinity of the PUM2 site (see also SI Materials and Methods and Fig. S4). Nevertheless, to eliminate this potential confounding factor and to facilitate the interpretation of structural effects, we focused only on variants without stable predicted structure in the protein-bound state (77% of variants).

The third condition is that the RBP must have known sequence specificity to pinpoint the RNA residues whose accessibility determines affinity and to account for all potential binding sites. Our recently developed thermodynamic model of PUM2 specificity allowed us to identify sequences in which the designed consensus or mutant sites were the only sites predicted to bind with significant affinity (SI Materials and Methods)(27). Alternative sites that are more accessible than the designed consensus site could make the structure effects appear weaker. The presence of additional sites would further allow for binding of multiple PUM2 monomers, and potential cooperativity, a feature that we address elsewhere (WRB et al., in revision). Herein, we excluded RNA variants that contained alternative binding sites outside the designed binding site for RNAs with sites having predicted affinities within the measurable affinity range (SI Materials and Methods).

We also assessed the possibility of PUM2 binding to partially accessible sites, which could decrease the observed structure effects. I.e., if a subset of the residues in the PUM2 site are substantially more accessible than the full site, and if loss of binding interactions due to the remaining occluded residues is less than the differential stability of the alternative structure, the observed effects of structure will be weaker than predicted for the full site. Calculations of predicted accessibilities of various partial sites using RNAfold and the corresponding predicted affinities using the thermodynamic PUM2 model suggest that (1) for the vast majority of variants (98.1% of UGUAUAUA variants), partial 4–7mer sites are not more accessible than full sites and (2) in cases where different accessibilities are predicted, the increase in accessibility for partial sites would not lead to enhanced binding (SI Materials and Methods).

A further test of the coupled binding model is that increasing temperature should have distinct effects on PUM2 binding to variants that contain structure vs. those that do not. Binding to unstructured variants is expected to become weaker at higher temperature, whereas for structured variants, destabilization of RNA structure should lead to increased accessibility of PUM2 sites, thereby counteracting the weaker intrinsic binding and resulting in smaller observed temperature effects. Indeed, this prediction is met, and, for the most stable structures, higher binding affinity is observed at 37 °C than at 25 °C (Fig. S5). As PUM2 met the criteria for application of the thermodynamic binding model of Fig. 1A and the effect of temperature was consistent with our expectation, we used it to quantify RNA accessibility due to structure and to test NN algorithms.

### Nearest-neighbor (NN) algorithms do not accurately predict RNA structure

To compare predicted and observed stabilities of RNA structure, we calculated the observed destabilization of PUM2 binding to each oligonucleotide relative to the unstructured state 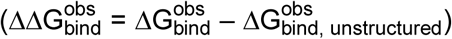 and plotted these values against secondary structure stabilities predicted by RNAfold (Turner 2004 parameters). We first assessed the results for PUM2 consensus sites at 25 °C, which provided the widest dynamic range of measured values. As expected, there was a decrease in binding affinity 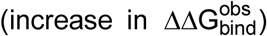 with increasing predicted secondary structure stability (Fig. 2A; R^2^ = 0.72). But despite this strong correlation, the observed effects of RNA structure were systematically and substantially lower than predicted, with a best fit slope of 0.47 rather than the slope of unity as predicted by the model (Fig. 2A), and with median deviations of up to ∼3 kcal/mol, corresponding to >200-fold overestimation of stability for RNAs with the greatest predicted stabilities (>4 kcal/mol) (Fig. 2B). Further, there was substantial variation in observed affinities for RNAs with similar predicted stabilities, as seen visually in Fig. 2A and as reflected in the correlation coefficient of R^2^ = 0.72 that is substantially less than that of 0.96 for replicate experiments (Fig. S3). Analogous results were observed for mutant PUM2 sites and for the experiment carried out with the PUM1 protein, a homolog of PUM2 with an identical consensus binding site (Fig. S6), providing strong independent support for these observations. Deviations of this magnitude indicate that this NN algorithm does not quantitatively predict the stability of RNA structures.

**Fig. 2.**
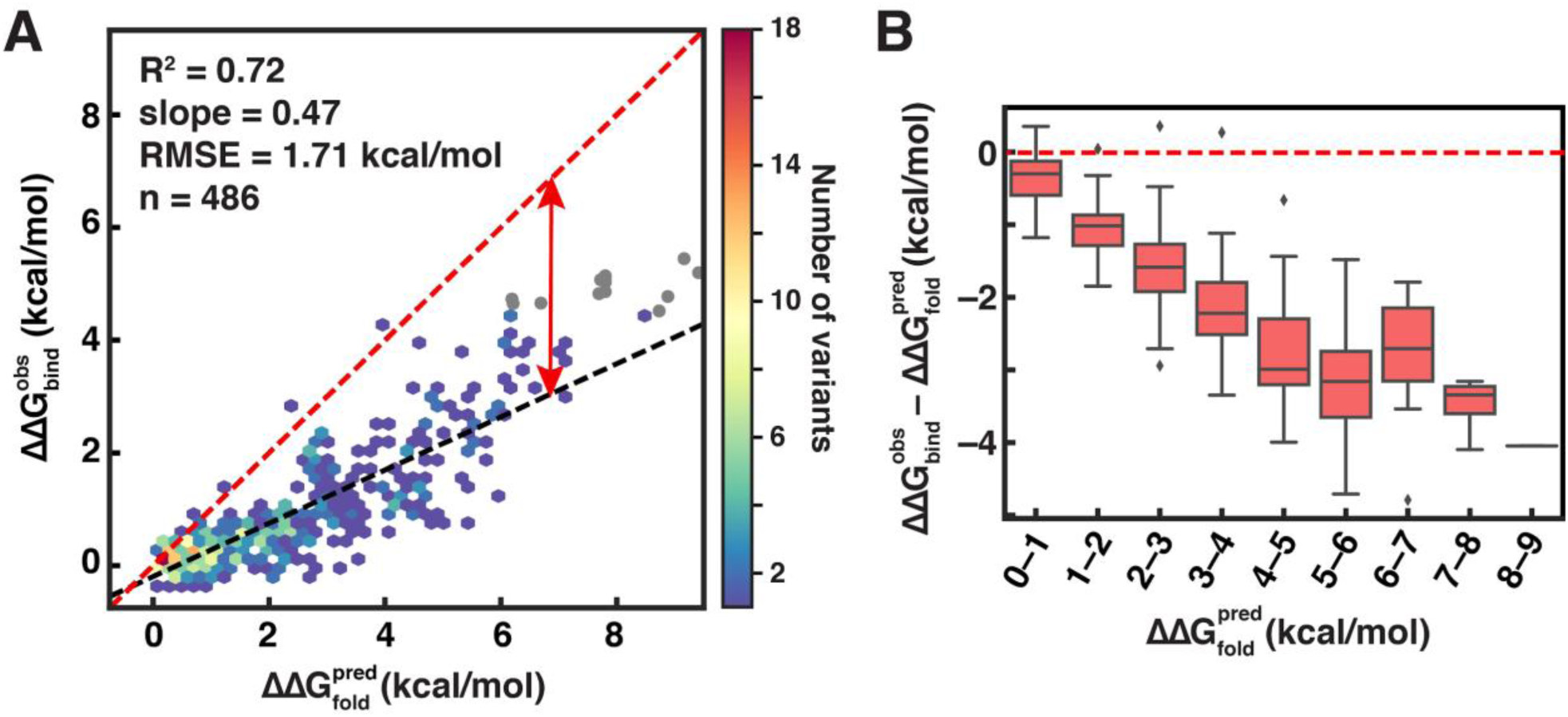
RNA structure weakens PUM2 binding in a stability-dependent fashion. (A) Scatterplot of predicted structure effects on binding 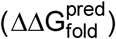 and observed weakened PUM2 binding 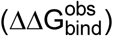 relative to UGUAUAUA sites predicted to lack structure. The predicted values are based on ensemble stabilities calculated using RNAfold with Turner 2004 parameters, and the measured values correspond to the weighted means of two replicate experiments at 25 °C (100 mM KOAc, 2 mM MgCl_2_, 20 mM Na-HEPES, pH 7.4, 0.1% Tween-20, 5% glycerol, 0.1 mg/ml BSA, and 2 mM DTT). Variants with measured affinities below the limit for high-confidence affinity determination are shown in gray and were not used in the correlation. The red dashed line indicates the predicted slope of 1, and the black dashed line indicates the best fit. (B) Differences between predicted and observed structure effects across predicted stability bins. Red line indicates agreement with the predicted values (corresponding to the diagonal in 2A).

Given the deviation between predicted and observed RNA stability for RNAfold, we asked whether other RNA prediction programs were able to better capture the observed energetics. Specifically, we tested: RNAstructure (11), UNAFold (originally Mfold) (30, 31), NUPACK (32), and additional parameter sets provided with the RNAfold installation. We found similar levels of agreement by all methods, with best fit slopes of 0.44 to 0.50 (Fig. 3A) and similar deviations from the predicted line of slope 1 (Fig. S7).

**Fig. 3.**
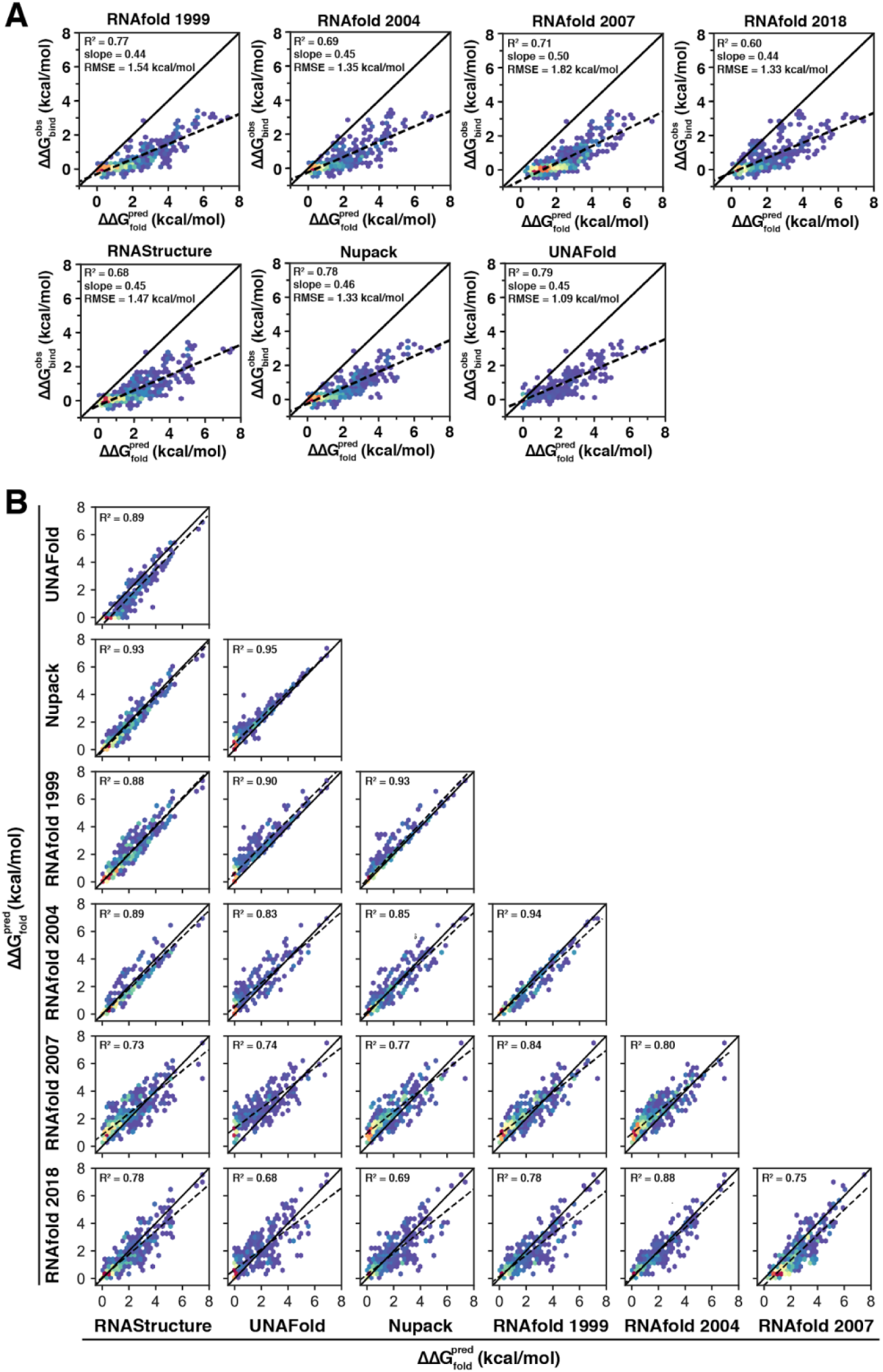
Comparison of secondary structure predictions based on various algorithms and parameter sets. (A) Comparisons of predicted and observed structure effects (37 °C). Data for UGUAUAUA-containing variants lacking predicted structure in the protein-bound state are shown (n = 620). Solid lines indicate perfect agreement between predicted and observed values; dashed lines indicate the fit slopes. To enable comparisons between all algorithms, predictions and data collected at 37 °C were used, because NUPACK and UNAfold do not provide up-to-date enthalpy parameters. (B) Pairwise comparisons between secondary structure predictions based on different algorithms and parameter sets. The same set of UGUAUAUA-containing oligonucleotides as shown in part *A* was used (37 °C; n = 620). See Table S2 for RMSE and slope values.

We also directly compared the predictions made by the different algorithms and parameter sets for our RNA library (Fig. 3B; Table S2). There are substantial quantitative differences between algorithms, even when based on the same parameter set (Table S2). There was less agreement between predicted and measured values than between algorithms (Fig. 3A,B), suggesting that there are stability factors not accounted for in any of the algorithms.

### Assessing the effects of ionic conditions on NN parameters

The underlying thermodynamic parameters of NN models originate from a single set of ‘standard’, high-salt conditions (1 M sodium chloride). We carried out experiments under ionic conditions (2 mM Mg^2+^, 100 mM K^+^, 8 mM Na^+^) closer to physiological (∼1 mM Mg^2+^, ∼150 mM K^+^), as these algorithms have been predominantly applied to evaluate the role of structure in cellular RNAs (33-36). Given the non-trivial relationship between RNA structural stability and salt conditions, extrapolation to different salt concentrations is currently not possible with RNA NN algorithms. Nevertheless, empirical relationships based on tightly bound ion theory between salt conditions and the stabilities of RNA hairpins have been developed (37, 38). We therefore applied these relationships to provide a rough estimate for the extent to which differences in salt conditions may contribute to the deviations from NN predictions observed in our experiments.

Applying this calculated salt adjustment generally weakened predicted structure stabilities by 0.7–1.3 kcal/mol for the RNAs that estimates could be derived for (Fig. 4A; SI Materials and Methods). Incorporating these corrections led to modest increases in the slope of the predicted vs. observed structural stabilities, but substantial deviations from the predicted slope of 1 still remained (Fig. 4B–D). This improvement suggests that a correction for salt is needed, and accounting for ionic conditions would likely improve predictions. Collection of empirical data under physiological salt conditions and continued development and testing of models of ionic effects on RNA stability are needed. Finally, there is evidence that RNA structure in cells is substantially less stable than predicted by NN algorithms (27). Some, but very likely not all, of this difference may be due to differences in physiologic ionic conditions and the ionic conditions used to obtain NN parameters.

**Fig. 4.**
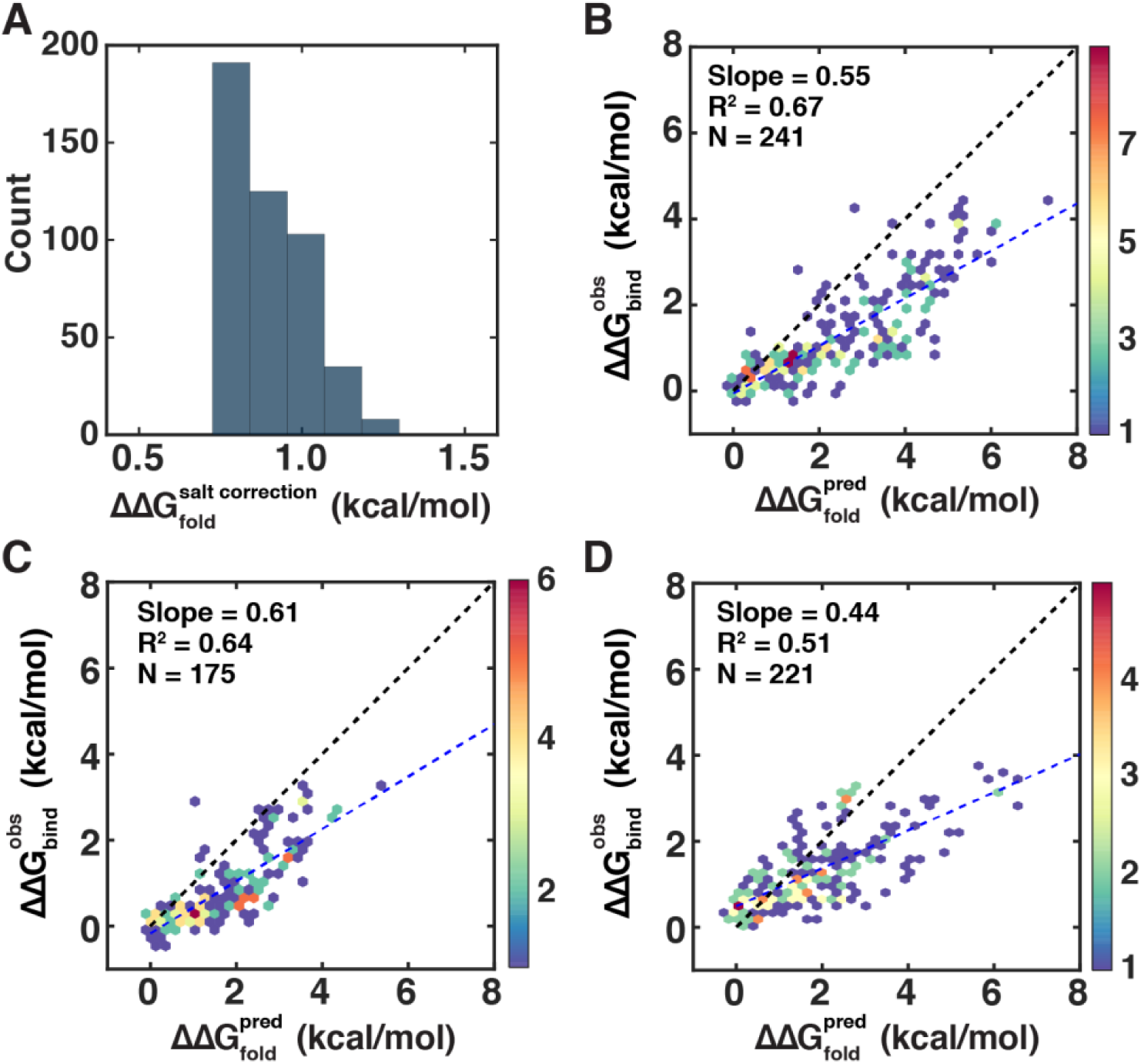
Salt-adjusted predictions of RNA secondary structure stability. (A) Histogram of the magnitude of the salt corrections based on the model from Tan and Chen (37, 38) for the RNA variants in this study. (B–D) Comparisons of predicted salt-adjusted structural stabilities to observed ΔΔG values for variants containing consensus (UGUAUAUA) sites at 25 °C (B), consensus (UGUAUAUA) sites at 37 °C (C), and mutant (UGUAUAUU) sites at 25 °C (D). Colors in B–D represent the density of points.

### Loop parameter inaccuracies account for major deviations

Given that the stabilities of the vast majority of RNA structural features have not yet been experimentally determined, incorrect parameter values used by NN algorithms seem likely to contribute to deviations. To begin to assess individual parameters, we designed our library to focus on a limited number of structural features in multiple sequence contexts with the majority of hairpins in our library designed to have UUCG and CCCC tetraloops. These loops were chosen as they exhibit average and exceptionally low tetraloop stability, respectively (39-43)^2^.

To assess the accuracy of predictions for each loop sequence, we divided our library into groups of RNAs with CCCC or UUCG loops, omitting from this analysis the se t of RNAs where neither of these two loops was predicted to be present in the most stable structure (Fig. 5A, B). The data clustered based on the loop sequence, with CCCC loops deviating from predicted values much more than UUCG loops (red (CCCC) and blue (UUCG) points in Fig. 5A, B). The distinct behaviors of constructs with these different loop sequences strongly suggests that parameter errors, rather than effects of experimental conditions––which would affect all variants––are the main source of the observed deviations.

**Fig. 5.**
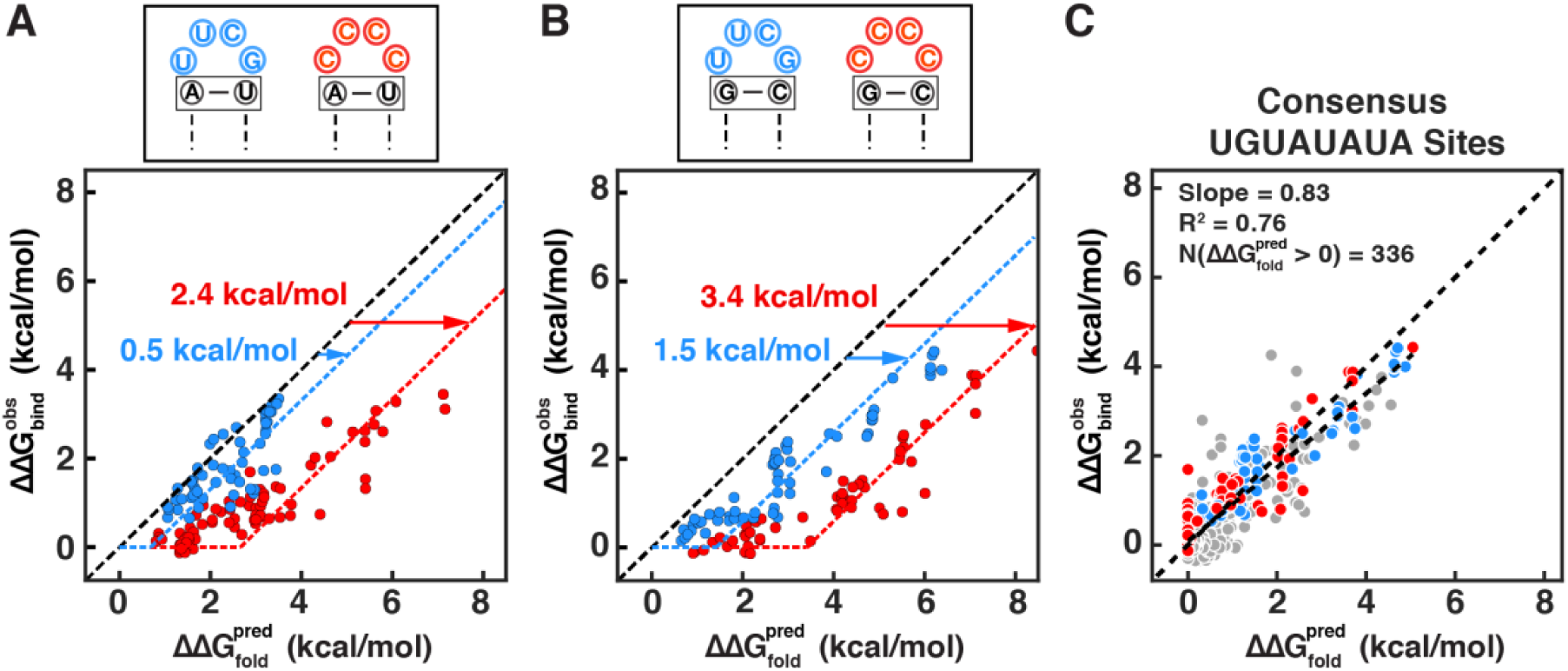
Prediction accuracy varies with the loop and closing base-pair sequence. (A,B) Correlation between predicted and observed structure effects for constructs containing UUCG (blue) vs. CCCC (red) loops in the minimum free energy structure predicted by RNAfold. The constructs were further divided based on the closing base-pair—A·U (A) or G·C (B)—in the predicted structure. Dashed lines indicate the fit of UUCG (blue) and CCCC (red) loop data to a model in which the loop initiation energy is offset by a constant amount (horizontal axis intercept), and then the data follows the expected model with slope = 1. The best-fit offset values suggest that the loops are less stable than predicted by the indicated amounts. See Fig. S7 for determination of the offset values. (C) Correlation of the predicted structural stability after correcting each loop type (UUCG, CCCC, or other) by constant offsets (from panel A and Fig. S8) vs. the experimentally observed effect on binding. Slope and R^2^ values were calculated based on the RNAs with 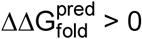.

The data with both types of loops exhibited a clear apparent ‘lag’ in plots of the predicted structural stability vs. observed binding affinity (Fig. 5A, B); up to a certain predicted stability, the structures containing a given loop showed no effect on binding, followed by an incline with a slope of approximately 1. A fraction of this offset likely reflects the effect of differences in salt conditions as described above, while the remainder must reflect differences between the loop initiation penalties used in RNAfold and the true loop initiation penalty. By fitting a model with a slope of 1 and different intercepts to the RNAs containing CCCC loops, UUCG loops, and other loops we were able to derive the total offsets for each of these loop types (Fig. 5A, B and Fig. S8). The fit offset for UUCG tetraloops of 0.7–1.4 kcal/mol (25 °C) is similar to the effects of ionic conditions estimated above and so could reflect ion effects rather than incorrect loop energy parameters (Fig. 5B, blue arrow). However, the larger offset for CCCC loops (2.7 kcal/mol or 3.4 kcal/mol with closing A·U or G·C base-pairs) suggests that differences in the loop initiation penalty from those used in RNAfold also contribute to the observed deviations (Fig. 5; 25 °C).

In principle, these values could be used to correct the values in RNAfold and better account for the observed data. As a simple demonstration, we applied the fit offsets representing both the salt correction and loop initiation penalty correction (Fig. 5A and 5B and Fig. S8) to RNAs containing each type of loop sequence (UUCG, CCCC, or other). As expected, this model gives a much-improved fit, with a slope of 0.81 instead 0.47 (Fig. 5C vs. Fig. 2). Extension of this approach to more structural features, via RNA-MaP or an analogous approach, could be used to obtain parameters for a more comprehensive RNA stability prediction algorithm.

With respect to the differences in the performance of RNA structure prediction algorithms for our data set (Fig. 3), these differences can be largely explained by differences in the loop penalties associated for UUCG and CCCC loops used in each algorithm (Table S3). For example, RNAfold does not account for the exceptionally low stabilities of oligo-C loops, despite their well-established properties that are included in the published parameter sets, leading to systematic underperformance for CCCC loop-containing hairpins (11, 29, 44). These systematic differences further underscore the value of independent tests.

## Concluding Remarks

RNA structure plays important biological roles, and NN algorithms provide a powerful framework for RNA structure predictions, representing the culmination of heroic efforts of multiple labs over decades. Nevertheless, despite considerable predictive power, our quantitative tests reveal that substantial deviations remain between prediction and experiment. We show that accounting for experimental salt conditions improves thermodynamic predictions, underscoring the importance of salt adjustments when applying NN algorithms, a feature important for drawing conclusions about cellular RNA structure and its biological roles. Additionally, substantial discrepancies between predicted and observed stabilities can be attributed to algorithm-specific differences in implementing known thermodynamic parameters. Further, the limited number of thermodynamic measurements obtained prevents determination of unique energies for each loop, thus requiring approximation of loop penalties based on averages from a small number of measurements. This approximation, given known substantial sequence-dependent variation (11), likely accounts for much of the remaining scatter between observed and predicted affinities in our experiments.

To test NN algorithms, we developed a quantitative, high-throughput assay based on protein binding, rather than UV melting, that provides advantages for the assessment of RNA secondary structure. Notably, this approach does not require extrapolation from higher temperatures and does not require a two-state assumption. This latter assumption will become increasingly limiting with RNA complexity, as RNAs are more likely to form ensembles of states and are more likely, within each state, to exhibit complex melting behavior. The method employed herein avoids these limitations by directly assessing ensemble stability. Thus, RNA-MaP and the protein-coupled structure assay provide and important tool for increasing the accuracy of RNA stability models.

In closing, we emphasize the general importance of blind tests of models and algorithms. Blind tests are a powerful means to assess and benchmark the quality of models, to identify shortcomings, and ultimately to develop more accurate predictive models.

## Materials and Methods

Full details are provided in SI Materials and Methods.

### Library design and preparation

The oligonucleotide library was ordered from CustomArray and is shown schematically in Fig. 1C and Fig. S1. The oligonucleotides were amplified using emulsion PCR (45) and PCR-assembled with constant adapter regions to form the RNA array construct depicted in Fig. 1B. Sequencing was performed using Illumina MiSeq Reagent Kit v3 (150-cycle).

### Protein expression, purification and labeling

Details of the preparation of SNAP-tagged RNA-binding domains of human PUM1 and PUM2 and labeling with Cy3B were described in (27). Active fractions of Cy3B-labeled protein, as determined by gel-shift titration experiments were as follows: 57% (PUM2-SNAP), 61% (SNAP-PUM1). Cy3B labeling efficiencies were: 60% (PUM2-SNAP), 53% (SNAP-PUM1).

### Binding experiments

RNA-MaP binding measurements were performed using a repurposed Illumina GAIIx instrument with a custom fluidics adapter interface, as described in (19, 21). Double-stranded DNA was regenerated on a sequenced Illumina MiSeq flowcell using biotinylated primers. Following incubation with streptavidin, RNA was generated using *E. coli* RNA polymerase. Stalling of the RNA polymerase at the terminal streptavidin block resulted in an array of RNA clusters, each containing ∼1000 transcripts. To determine PUM1 and PUM2 affinities, Cy3B-labeled PUM1 or PUM2 were flowed at increasing concentrations in Binding buffer (20 mM Na-HEPES, pH 7.4, 100 mM KOAc, 0.1% Tween-20, 5% glycerol, 0.1 mg/ml BSA, 2 mM MgCl_2_ and 2 mM DTT). Following equilibration, bound protein was imaged with a green laser (530 nm with a 590 nm band pass filter; 400 ms at 200 mW). Protein solutions were incubated for times ranging from 33 min for the lowest concentrations to 19 min for the highest protein concentrations at 25 °C, and for 15–23 min at 37 °C.

The fluorescence data were quantified as described in (21, 27). The fluorescence values at each protein concentration, normalized by the amount of transcribed RNA per cluster, were fit to a single-site equilibrium binding equation with a non-specific binding term (27):

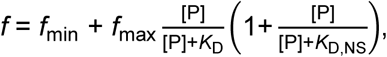

where f_min_ is the background fluorescence, f_max_ is the fluorescence at saturation, [P] is the concentration of the protein in solution (here, [P] ≈ [P]_total_), *K*_D_ is the dissociation constant (*K*_D_ = e^ΔG/RT^) and *K*_D,NS_ is the non-specific dissociation constant for a second protein monomer that binds to the RNA/protein complex 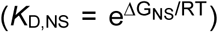. This model was used to account for an observed increase in fluorescence at high concentrations of protein after apparent saturation (Fig. S2).

### RNA structure predictions

Initial predictions were performed using Vienna RNAfold (v2.4.10) with the Turner 2004 parameter set (‘rna_turner_2004’)(11, 29). Additional parameter sets available with the Vienna 2.4.10 installation were also tested: rna_turner1999 (44), rna_andronescu2007 (46), and rna_langdon2018 (47). To compare secondary structure predictions across different algorithms, RNAstructure v6.0.1 (11), NUPACK v3.0 (32), and UNAFold v3.8 (31) were used, with details provided in SI Materials and Methods.

### Calculating salt-adjusted stabilities

Salt adjusted stabilities were calculated using empirical relationships derived by Tan and Chen (37, 38). The MFE structure predicted by RNAFold was used as the reference for identification of the lengths of hairpin stems, hairpin loops, and internal bulges. Salt contribution was computed at 1M NaCl and at the conditions used for our experimental measurements (108 mM monovalent and 2 mM Mg^2+^). The difference was then used to determine the salt adjusted 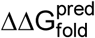.

### Fitting loop penalties

The free energy offsets for each loop type were determined by finding the intercept resulting in the lowest RMSE value for a line with slope 1. The RMSE value for each intercept was calculated using only the points with 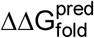 less than the current intercept value (i.e., all points were used for an intercept of 0 and a decreasing number of points were considered as the intercept increased).

## Acknowledgements

This work was funded by grants from the U.S. National Institutes of Health: P01 GM066275 (D.H., R.D., W.J.G.), R35 GM122579 (R.D.), R01 GM121487 and R01 GM111990 to (W.J.G), and by the Beckman Center. W.J.G acknowledges support as a Chan-Zuckerberg Investigator.

## Footnotes

1 The number includes four possible WC closing base pairs.

2 UUCG tetraloops are known to have exceptionally high stability in the presence of a C·G closing base-pair, but not with the closing base-pairs used in this study (41).

