## Supplemental Text and Figures for "Quantitative high-throughput tests of ubiquitous RNA secondary structure prediction algorithms via RNA/protein binding"

### SI Materials and Methods

#### Library design

The oligonucleotide library was designed to contain PUM1/2 protein binding sites within RNA hairpins of varying stability. The following features were varied, individually or combinatorially: the length and the sequence of the hairpin stem; the length and the sequence of the hairpin loop; the number and position of mismatches in the hairpin stem; the presence and size of bulges in the hairpin stem; the sequences flanking the hairpin. In each category, described below, we also varied the location of the PUM1/2 site (i.e., within the 5' or 3' strand of the hairpin), as well as the identity of the PUM1/2 site (UGUAUAUA consensus or UGUUAUAU mutant).

The individual sublibraries are shown in Fig. S1 and are described below, with numbering matching that in Fig. S1:

##### 1) Variable length and sequence of a continuously base-paired stem region.

- A series of 1–10 base-pairs (for the consensus site) or 1–9 base-pairs (for the mutant site) was introduced in every register within the region indicated in Fig. S1, section 1. Here and below base-pairs refer to canonical Watson-Crick base-pairs (WC bps).
- All terminal loops contained either the 'UUCG' or 'CCCC' sequence (in constructs in which the base-paired region was located one or more nucleotides away from the indicated loop, longer loops resulted due to additional non-basepaired residues flanking the UUCG or CCCC sequences).
- The 5' and 3' sequences flanking the hairpin consisted of either 'UUU' triplets at each terminus, 'AAA' triplets at each terminus or a combination of 'UUU' and 'UCUCUU' sequences, with the 'UCUCUU' sequence positioned at the terminus near the PUM1/2 site. The 'UUU'/'UCUCUU' flanking sequences were used throughout the other sublibraries. No systematic effects of flanking sequences were observed on the correlation between predicted and observed structure effects.
- Here and below, unless otherwise indicated, the PUM1/2 consensus site was consistently preceded by 'CU' residues to enable application of this library to the homologous yeast protein Puf3, which contains an additional binding pocket for an upstream C residue. Similarly, in the case of the mutated site, the UGUUAUAU sequence was consistently followed by an A residue, to enable applications to the yeast Puf4 protein, which binds a 9-nt site with the residue at position 7 preferentially flipped out.

##### 2) Variable length and sequence of the hairpin loop.

- The variants contained either no added terminal loops (0 nt) or contained loops of the following lengths, with two C/U-containing sequences designed per loop length: 1, 2, 3, 4, 5, 6, 8, 10, 15, 20, 30, 40, 50 nt.
- The hairpin stems consisted of either a perfectly complementary 8-bp region or included one to two mismatches.

3) Single mismatches.

- Single mismatches were introduced at varying positions of a 5–8 bp long stem regions (4–7 bp final), in the presence of UUCG or CCCC tetraloops.

4) Double mismatches.

- Two mismatches, adjacent or separated by one or more base-pairs were introduced into 6–8 bp long stem regions (4–6 bp final), in the presence of UUCG or CCCC tetraloops.

5) Bulges opposite the PUM1/2 site.

- Insertions of 1–4 nt were introduced into the strand complementary to the PUM1/2 site, in the presence of 5–8 bps and UUCG or CCCC tetraloops.

6) Bulges within the PUM1/2 site.

- The hairpin strand opposite the PUM1/2 binding site was designed to leave 1–6 residues in the PUM1/2 site bulged out. The number of bps flanking the bulge was also varied, and the terminal loops were UUCG or CCCC.

The sequences for each structure category were designed using custom Python functions and were curated to prioritize variants that were predicted to form the intended structure in RNAfold (1). Nevertheless, when designing the library, we did not strictly require that all sequences form the designed structure in their predicted minimum-free energy (MFE) state, given the limitations of structure prediction algorithms, the heterogeneity of RNA structural ensembles, and our goal of broadly testing the accuracy of nearest-neighbor stability predictions. The MFE structures predicted by RNAfold for all variants are indicated in the dot-bracket notation in Suppl. Dataset 1.

An overarching consideration in designing the library was that the magnitude of predicted structural stabilities should not greatly exceed the measurable PUM1/2 affinity range, as structures with greater stabilities would completely ablate binding, preventing quantitative analysis. Correspondingly, the library consisted of short hairpin constructs containing no more than 10 bp. For PUM2 binding at 25 °C, the range of affinities that could be quantified with high confidence was 4.7 kcal/mol. Because of generally weaker-than-predicted structure effects, we were able to cover a range of predicted stabilities of 8.5 kcal/mol, with more stable structures giving no detectable binding (Fig. 2, Fig. S2).

#### **Library preparation and sequencing**

The preparation of a larger library that contained the structured oligonucleotides described above was fully described previously in (2). Briefly, DNA oligonucleotides were ordered from CustomArray, amplified using emulsion PCR (3) and PCR-assembled with constant adapter regions to form the RNA array construct schematically depicted in Fig. 1B. The library was bottlenecked to ensure that multiple copies of each barcode were represented on the final RNA array, and sequencing was performed using Illumina MiSeq Reagent Kit v3 (150-cycle; 56 nt in Read 1, 96 nt in Read 2). To ensure proper separation of RNA clusters during subsequent binding experiments, the oligonucleotide library was combined with an excess of

PhiX DNA (Illumina; 84–90% of the total 6–9.6 fmol DNA). Additionally, a fiduciary marker oligonucleotide was spiked in at 1% and was used for image alignment during binding experiments (2).

#### **Protein expression, purification and labeling**

Details of the preparation of SNAP-tagged RNA-binding domains of human PUM1 and PUM2 and labeling with Cy3B were fully described in (2). Active fractions of Cy3B-labeled protein, as determined by gel-shift titration experiments were as follows: 57% (PUM2-SNAP), 61% (SNAP-PUM1). Cy3B labeling efficiencies were: 60% (PUM2-SNAP), 53% (SNAP-PUM1).

#### **Binding experiments**

RNA-MaP binding measurements were performed on a larger library of 25,052 oligonucleotides and were described in detail in (2). Briefly, a repurposed Illumina GAlIx instrument with a custom fluidics adapter interface was used, as described in (4, 5). An Illumina MiSeq flowcell containing the sequenced oligonucleotide library was washed, and double-stranded DNA was generated using Klenow fragment (3'-5' exo(-); NEB) with a biotinylated primer. Streptavidin was then added to generate a block at the end of the DNA template for stalling and immobilizing the RNA polymerase in the subsequent transcription step. RNA was generated using *E. coli* RNA polymerase holoenzyme (NEB) at 37 °C. To ensure loading of a single RNA polymerase molecule per DNA template, RNA polymerase was first incubated with Initiation buffer, which lacked CTP (20 mM Tris•HCl pH 8.0, 7 mM MgCl<sub>2</sub>, 20 mM NaCl, 0.1% BME, 0.1 mM Na<sub>2</sub>EDTA, 1.5% glycerol, 0.02 mg/mL BSA, 0.01% Tween 20, and 2.5 μM each of ATP, GTP, and UTP). This resulted in stalling of the enzyme upon encountering the first cytosine in the 'RNA polymerase initiation and stall' region (Fig. 1B). After washing away excess RNA polymerase, transcription was re-initiated in Extension buffer, which contained all 4 ribonucleotides (20 mM Tris•HCl pH 8.0, 7 mM MgCl<sub>2</sub>, 20 mM NaCl, 0.1% BME, 0.1 mM EDTA, 1.5% glycerol, 0.02 mg/mL BSA, 0.01% Tween 20, and 1 mM each of ATP, GTP, UTP and CTP). During this step, oligonucleotides complementary to the constant sequences flanking the variable region in the RNA transcripts were also added to prevent undesired intramolecular interactions. The oligonucleotide complementary to the 5' end of the transcript was 3'-labeled with Alexa Fluor 647 (IDT) to enable the visualization and quantification of the transcripts (Fig. 1B). Transcription was followed by an additional incubation with the blocking oligonucleotides. The flow cell was subsequently washed with Binding buffer (20 mM Na-HEPES, pH 7.4, 100 mM KOAc, 0.1% Tween-20, 5% glycerol, 0.1 mg/ml BSA, 2 mM MgCl<sub>2</sub> and 2 mM DTT), the temperature was adjusted to 25 °C or maintained at 37 °C. The flow cell was imaged in the red channel (660 nm laser with a 664 nm long pass filter; 400 ms at 200 mW) to visualize the transcript-annealed fluorescent probes, and in the green channel (530 nm laser with a 590 nm band pass filter; 400 ms at 200 mW) to visualize background and the fiducial markers (visualized with 5' TYE 563-labeled complementary oligonucleotide (IDT)) (2).

To determine PUM1 and PUM2 affinities, Cy3B-labeled PUM1 or PUM2 were flowed at increasing concentrations in Binding buffer, equilibrated and imaged in the green channel. Protein solutions were incubated for times ranging from 33 min for the lowest concentrations to 19 min for the highest protein concentrations at 25 °C, and for 15–23 min at 37 °C. Sufficient equilibration time was determined based on the following: 1) Dissociation time courses, measured for PUM2 by rapidly flowing 3 μM of consensus-containing oligonucleotide (UCUUGUAUAUAUA) with simultaneous imaging; 2) Consistent amplitudes of binding curves irrespective of predicted structure stability, which indicated lack of accumulation of stable structures that do not equilibrate on the time scale of the experiment (Fig. S9); 3) previous gel-shift measurements of PUM2 dissociation from consensus RNA at 25 °C indicated rapid equilibration (halflife of 6.1 min; (6)), setting the upper limit for equilibration at 30.5 min (five halflives).

#### Data fitting

The fluorescence data were quantified as described in (2, 5). To account for variable amount of transcribed RNA per cluster, fluorescence in the green channel (protein colocalization) was normalized by the fluorescence in the red channel (amount of transcribed RNA). The normalized fluorescence values at each protein concentration were fit to a single-site equilibrium binding equation with a non-specific binding term (2):

$$f = f_{\min} + f_{\max} \frac{[P]}{[P] + K_D} \left( 1 + \frac{[P]}{[P] + K_{D,NS}} \right) \quad (\text{Eq. 1}),$$

where $f_{\min}$ is the background fluorescence, $f_{\max}$ is the fluorescence at saturation, $[P]$ is the concentration of the protein in solution (here, $[P] \approx [P]_{\text{total}}$), $K_D$ is the dissociation constant ($K_D = e^{\Delta G/RT}$) and $K_{D,NS}$ is the non-specific dissociation constant for a second protein monomer that binds to the RNA/protein complex ($K_{D,NS} = e^{\Delta G_{NS}/RT}$). This model was used to account for an observed increase in fluorescence at high concentrations of protein after apparent saturation (Fig. S2).

#### Data filtering

Unless otherwise indicated, the high-confidence variants used in the analyses throughout the manuscript were determined based on the following criteria:

(1) the variant was represented by at least five clusters; for PUM2 replicate experiments at 25 °C, this condition had to be met in at least one replicate experiment. In case only one replicate had five or more clusters, the $\Delta G$ value from this replicate was used instead of the replicate average.

(2) 95% bootstrap confidence interval of the $\Delta G$ value was less than 1 kcal/mol; in the case of PUM2 replicate experiments, 1 kcal/mol cutoff was applied to the weighted replicate error.

(3) observed $\Delta G$ values were below $-8.5$ kcal/mol, which could be confidently distinguished from background, non-transcribed clusters (2) (this filter was applied before adjusting for active protein concentration).

(4) the variant lacked alternative sites with predicted affinities within the measurable range (see Identifying alternative binding sites below).

(5) the variant lacked predicted structure in the protein-bound state, as determined by constraining the 8mer PUM1/2 binding site to the single-stranded state in and calculating the stability of the resulting ensemble using RNAfold (see RNA structure predictions below).

Variants with $\Delta G_{\text{fold}}$ values for the constrained ensemble lower than $-0.5$ kcal/mol were excluded from the analysis.

Table S4 indicates the fractions of variants that meet the above criteria.

#### Identifying alternative binding sites

To ensure that the observed affinity effects reflected the accessibility of the designed site (consensus or mutant), rather than PUM1/2 binding to alternative, lower affinity sites, and to focus on variants with a single specifically bound PUM1/2 monomer, we removed variants that contained additional predicted binding sites outside the designed site. To identify these variants, we used the previously developed thermodynamic model for PUM1/2 binding to calculate predicted affinities for every 9mer, 10mer and 11mer binding register within each oligonucleotide—corresponding to sites with no flipped residues, one flipped residue, and two flipped residues, respectively (2). (These calculations accounted for weak position 9 binding, in addition to the core 8mer binding site; however, position 9 was not considered in determining overlap.) Variants that contained additional binding registers that did not overlap the designed binding site, and that had predicted $\Delta\Delta G$ values lower than 5 kcal/mol (25°C) or lower than 4.5 kcal/mol (37°C) were excluded from further analysis, ensuring no detectable binding within our measurable range. Variants containing more than one UGUA sites were also excluded.

#### RNA structure predictions

*RNAfold*. Initial predictions were performed using Vienna RNAfold (v2.4.10) with the Turner 2004 parameter set, corresponding to the most recent parameter set based on thermodynamic measurements, which is used as the default option in RNAfold (1, 7). The stability of the structural ensemble was determined with the following command: `RNAfold -p0 -T 25` or `RNAfold -p0 -T 37`, where the `p0` argument is used to calculate the ensemble free energy and `T` is the temperature in degrees Celsius.

To compare the predicted stabilities between the different parameter sets included in the Vienna package, the `-P` argument was additionally provided (e.g., `RNAfold -p0 -T 37 -P rna_turner1999.par` for Turner 1999 parameters (8)). Each of the four parameter sets available from the 'misc' folder of the Vienna 2.4.10 installation was tested: `rna_turner1999` (8), `rna_turner2004` (7), `rna_andronescu2007` (9), `rna_langdon2018` (10).

To predict the stability of the structural ensemble in which the PUM1/2 binding site is accessible, we ran the above command with a constraint argument (`-C`) and constrained the binding site (UGUAUAUA or UGUAUAUU) to non-base paired state. E.g.:

```
UCUCUUUGUAUAUAUCUCUU
.....xxxxxxxxx.....
```

*RNAstructure*. RNAstructure v6.0.1 (7) `partition` and `EnsembleEnergy` functions were used to calculate ensemble free energies at 37 °C: `partition <input.txt>`

```
<output_partition_file.txt> --temperature 310.15 and EnsembleEnergy
<output_partition_file.txt>.
```

**NUPACK.** NUPACK v3.0 (11) pfunc function was used with the ‘rna1999’ parameters (8): `pfunc -material rna1999 <input_seq>`.

**UNAFold.** The UNAFold.pl script from the UNAFold v3.8 installation (12) was used to calculate ensemble stabilities using the default Turner 1999 parameters (8): `UNAFold.pl <input.txt>`.

Comparisons of all prediction algorithms are indicated in Table S2. The ensemble stabilities predicted for our oligonucleotide library by each algorithm at 37 °C are indicated in Suppl. Dataset 1.

#### Comparing predicted vs. observed structure effects

The observed structure effects were calculated as the difference between the observed free energy of binding for a given variant and the median measured $\Delta G$ value for the unstructured instances of the consensus and mutant sequences, with unstructured sequences identified as having predicted folding free energy greater than $-0.2$ kcal/mol (RNAfold; Turner 2004 parameters):

$$\Delta\Delta G_{\text{bind}}^{\text{obs}} = \Delta G_{\text{bind}}^{\text{obs}} - \Delta G_{\text{bind, unstructured}}^{\text{obs}} \quad (\text{Eq.2}).$$

The unstructured variants included oligonucleotides from other sublibraries that were measured in parallel (2), with the full sequences and affinity information for these variants provided in Suppl. Dataset 2.

Our recent results revealed weak specificity of PUM1/2 for a G residue immediately following the PUM1/2 8mer binding site (position 9). To account for variation at this position in our structured oligonucleotide library for the consensus sites, we used the previously determined position 9 penalties to correct the $\Delta\Delta G_{\text{bind}}^{\text{obs}}$ value (2) (mutant sites were always followed by a U residue, see Library design). Variants containing a U residue at the 9th position were used as the reference (with unstructured UGUUAUAUAU variants used to calculate the median $\Delta\Delta G_{\text{bind}}^{\text{obs}}$ for the consensus sites), and the $\Delta\Delta G$ adjustments for the other bases were 0.30 kcal/mol (9A), 0.29 kcal/mol (9C) and $-0.07$ kcal/mol (9G) to account for the contribution of position 9 identity to the observed $\Delta G$ values:

$$\Delta\Delta G_{\text{bind}}^{\text{obs}} = \Delta G_{\text{bind}}^{\text{obs}} - \Delta G_{\text{bind, unstructured UGUUAUAUAU}}^{\text{obs}} - \Delta\Delta G_{9A/C/G} \quad (\text{Eq.3}).$$

The predicted structure effects $\Delta\Delta G_{\text{fold}}^{\text{pred}}$ were based on the ensemble $\Delta G$ values predicted by each of the algorithms: $\Delta\Delta G_{\text{fold}}^{\text{pred}} = -\Delta G_{\text{fold}}^{\text{pred}}$. To facilitate the analysis and interpretation of observed structure effects, variants with predicted significant structure in the

protein-bound state were not included in the analysis (Fig. 1A). All prediction programs except UNAFold limited the $\Delta G_{\text{fold}}^{\text{pred}}$ values to no greater than 0 kcal/mol; for consistency, and given that positive $\Delta G_{\text{fold}}^{\text{pred}}$ would not enhance binding, positive UNAFold values were replaced with 0 kcal/mol.

#### Assessing the effects of structures outside the PUM1/2 site

To evaluate the effects of RNA structure outside the PUM1/2 binding site on PUM2 binding, we compared the observed binding affinities ($\Delta G_{\text{bind}}^{\text{obs}}$ values) for all variants that contained an accessible PUM1/2 site and that were ($n = 24$) or were not ($n = 92$) predicted to contain structure outside the PUM1/2 binding site. To identify all variants with accessible PUM1/2 consensus sites (with or without adjacent structure), we first calculated the ensemble stabilities for each oligonucleotide using RNAfold, with and without constraining the PUM1/2 consensus site to the single-stranded state (1) (see **RNA structure predictions**). We then selected variants in which the predicted stabilities with and without constraints differed by less than 0.5 kcal/mol, indicating lack of significant predicted structure over the PUM1/2 binding site ($\Delta G_{\text{fold, constr.}}^{\text{pred}} - \Delta G_{\text{fold, unconstr.}}^{\text{pred}} < 0.5$ kcal/mol). Among these variants, variants that contained significant predicted structure outside the PUM1/2 site were defined as those having predicted ensemble folding free energy of the constrained ensemble below $-0.5$ kcal/mol ($\Delta G_{\text{fold, constr.}}^{\text{pred}} < -0.5$ kcal/mol).

#### Evaluating partially occluded sites

To assess potential contributions of partially accessible binding sites to the observed binding, first, we compared predicted accessibilities of complete (8mer) and partial (4–7mer) binding sites. Only partial sites expected to be bound within or close to our measurable range were included in the analysis. Affinities of partial sites were estimated based on the previously determined single mutant penalties (2), assuming additive contributions of individual positions and assuming that the effect of not forming the interaction at a given position corresponded to the average of single mutant effects at that position. To compare the accessibilities of the 4–7mer sites (UGUA, UGU AU, UGU AU A, UGU AU AU, and GU AU AU A) to 8mer accessibility, we used RNAfold to calculate the structural stabilities of ensembles in which 4–7mer sites were constrained to be single-stranded (see **RNA structure predictions**). Variants were curated as described in the **Data filtering** section. In 98.1% of cases, the predicted stabilities were within 0.5 kcal/mol regardless of the size of the constrained region, indicating that partial sites were not more accessible. In the remaining cases ($n = 8$), the greatest increases in accessibility, of up to 3.85 kcal/mol were predicted for UGUA and UGU AU sites. However, this predicted increase in accessibility is smaller than the predicted weakening of binding due to the loss of 3' interactions ($\Delta \Delta G_{\text{bind}}^{\text{pred}} = 4.72$ kcal/mol, based on average mutation penalties for positions 6–8), indicating that binding of partial PUM1/2 sites is not energetically more favorable even when they are more structurally accessible and that binding of such sites should not significantly contribute to the observed affinity.

#### Calculating salt-adjusted stabilities

Salt adjusted stabilities were calculated using empirical relationships derived by Tan and Chen (13, 14). Correcting for different salt conditions requires identification of the length of hairpin stems, hairpin loops, and internal bulges. To determine these lengths, the MFE structure was first identified with RNAFold. Following identification of the most stable structure and the corresponding structural features, we computed the salt contribution at 1M NaCl and at the conditions used for our experimental measurements (108 mM monovalent and 2 mM Mg<sup>2+</sup>). The difference was then used to determine the salt adjusted $\Delta\Delta G_{\text{fold}}^{\text{pred}}$. These corrections were only possible for a subset of our constructs as the empirical equations do not apply to RNAs with stem lengths $\leq 4$ bp. While the reliance of this analysis on the predicted MFE structure presents a limitation, it is likely not a major one due to the relatively constant correction calculated for all structures in our RNA library.

#### **Fitting loop penalties**

The free energy offsets for each loop type were determined by finding the intercept resulting in the lowest RMSE value for a line with slope 1. The RMSE value for each intercept was calculated using only the points with $\Delta\Delta G_{\text{fold}}^{\text{pred}}$ less than the current intercept value (i.e., all points were used for an intercept of 0 and a decreasing number of points were considered as the intercept increased). These fits and the sensitivities of the RMSE values to the different fit offsets are depicted in Fig. S8. For all loop fitting, variants with $\Delta G_{\text{fold}}^{\text{pred}}$ values for the constrained ensemble lower than $-0.2$ kcal/mol were excluded from the analysis. The offsets applied in Figure 5C come from fitting the loop penalties for UUCG and CCCC tetraloops with GC and AU closing base pairs and all other loops with any closing base pair across variants containing UGUUAUA binding sites.

Supplemental Figures

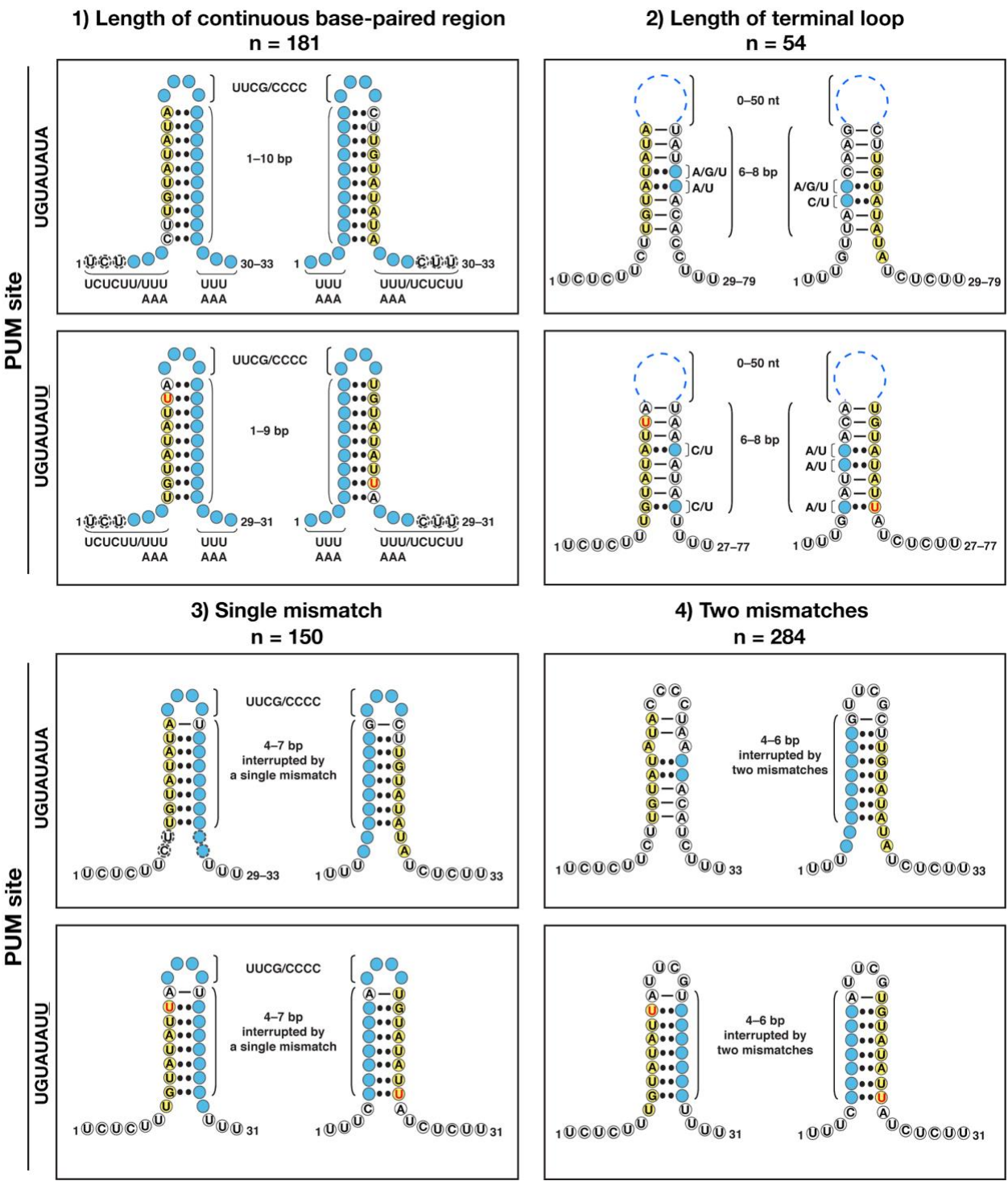

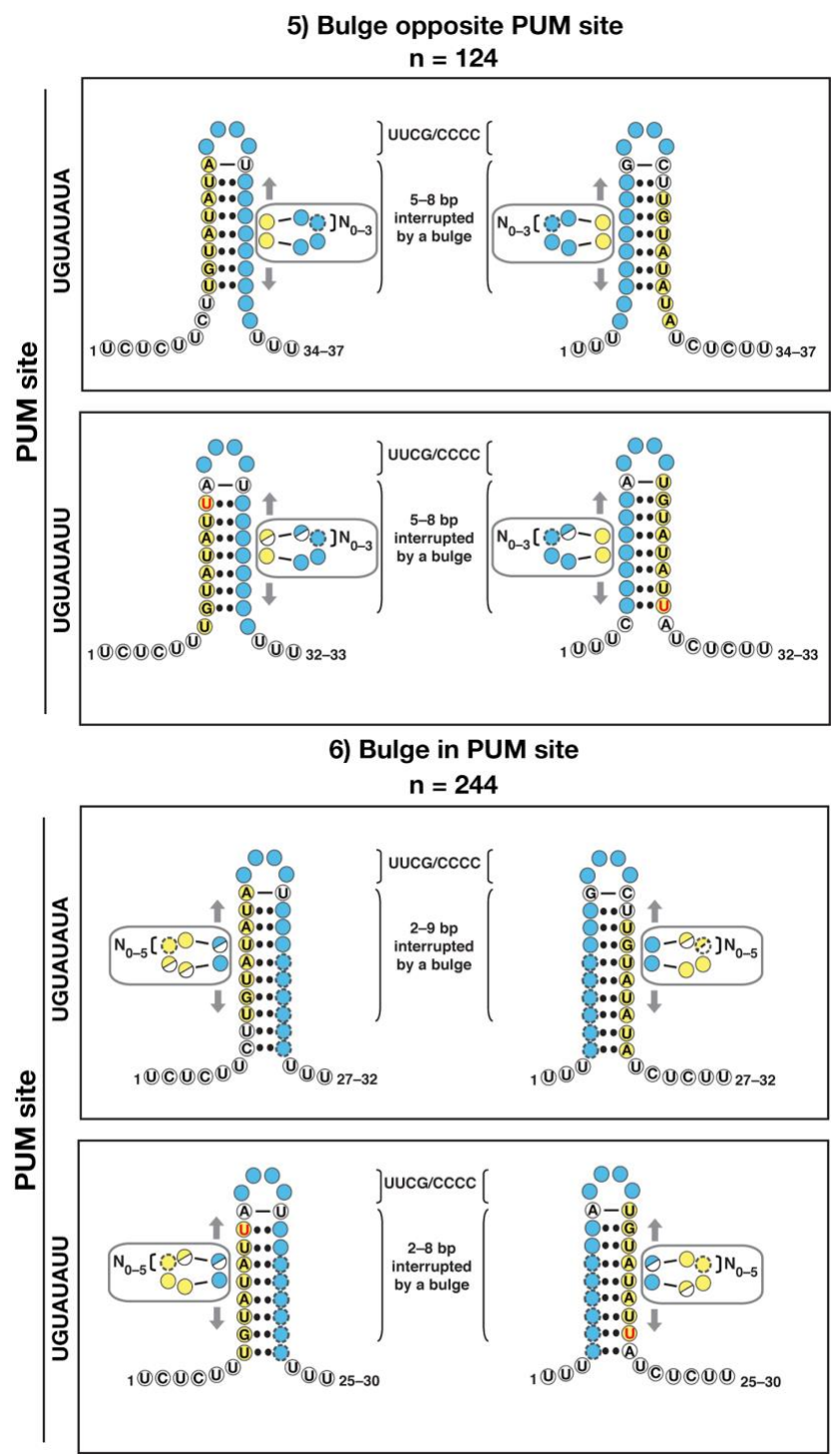

Fig. S1. Oligonucleotide library design for testing nearest-neighbor predictions via PUM1/2 binding. Each box includes designs in which the PUM1/2 site (yellow circles) is located in the 5′ or 3′ portion of the oligonucleotide. Each category includes constructs containing the PUM1/2 consensus (top) or mutated site (bottom), with the mutated nucleotide shown in red. ‘N’ indicates the number of variants for which high-quality PUM2 binding data were obtained at 25 ˚C. Watson-Crick (WC) base pairs that are always present at a given position are indicated with lines, and base pairs present only in a subset of constructs are indicated with two dots. Circles indicate individual nucleotides, with the color scheme as follows: *white*, positions with invariable sequence; *blue*, positions with variable sequence; *yellow*, PUM1/2 binding site. In parts 5 and 6, two-colored circles are used to indicate that, depending on the construct sequence and the position of the bulge along the stem, the bulges and surrounding base-pairs may include variable or invariant positions. Circles with dashed outlines indicate nucleotides that are present only in a subset of constructs. WC base-pairs are referred to as 'bp'.

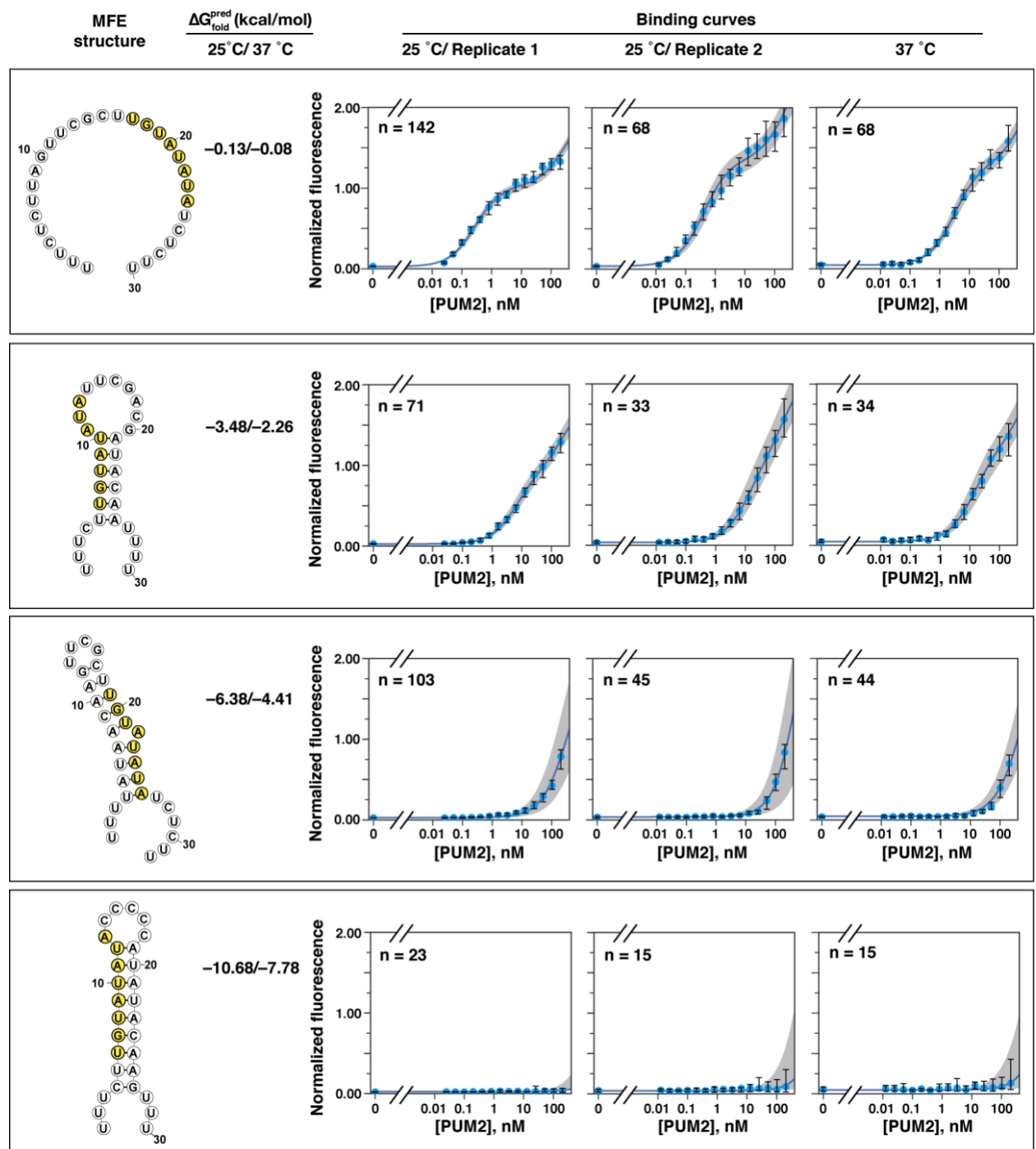

**Fig. S2.** Representative curves for PUM2 binding sites of different predicted stabilities. The slight increase in fluorescence at high concentrations for some of the weakest binders (bottom row) is beyond the limit at which binding can be confidently distinguished from background of non-transcribed clusters (2). The structure models of the minimum-free energy (MFE) structures (left) were generated by RNAfold, which was also used to calculate the stabilities of the folded ensembles ($\Delta G_{\text{fold}}^{\text{pred}}$; see Methods) (1). The PUM1/2 consensus site is highlighted in yellow.

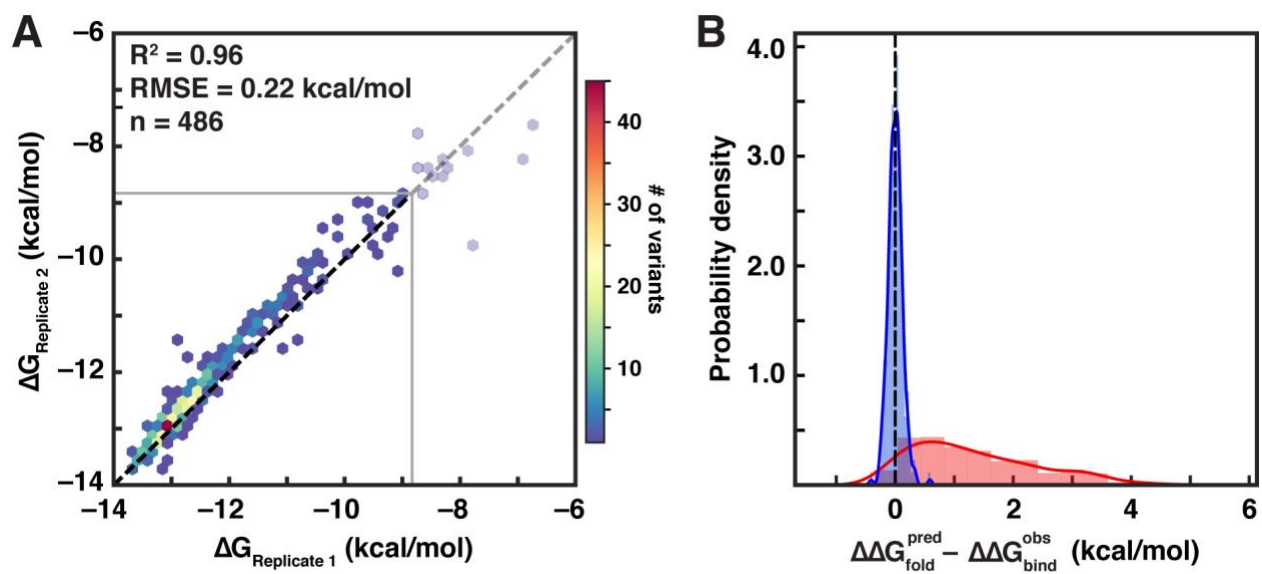

**Fig. S3.** Reproducibility of RNA-MaP measurements and comparison to prediction errors demonstrating that prediction error exceeds experimental errors. (A) Reproducibility between replicate experiments measuring PUM2 binding at 25 °C. The same variants as displayed in Figure 2A are shown. The RMSE value takes into account the constant offset of 0.28 kcal/mol between replicates, presumably due to differences in the active protein concentration. (B) Comparison of the distribution of weighted replicate errors (blue) vs. the differences between predicted and observed structure effects (red), which shows that the errors between the predicted and observed structural stabilities are much greater than would be expected if they were due to experimental error.

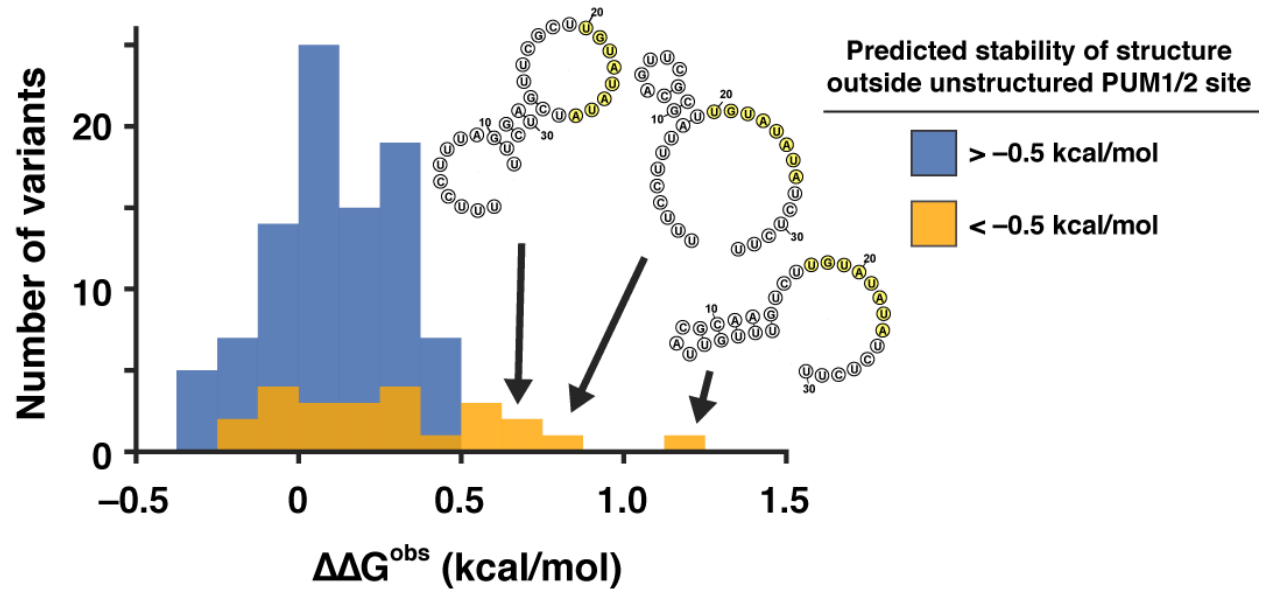

**Fig. S4.** Effects of predicted structure outside the PUM1/2 binding site on observed binding. To assess if RNA structure outside the PUM1/2 binding site affected PUM2 binding, we compared the observed binding affinities ($\Delta\Delta G_{\text{bind}}^{\text{obs}}$ values) for all variants that contained a single-stranded PUM1/2 site and that were (orange; $n = 24$) or were not (blue; $n = 92$) predicted to contain structure outside the PUM1/2 binding site. To identify all variants with unstructured PUM1/2 sites, we first calculated the ensemble stabilities for each oligonucleotide using RNAfold, with and without constraining the PUM1/2 consensus site to the single-stranded state (1). We then selected variants in which the stabilities with and without constraints differed by less than 0.5 kcal/mol, indicating lack of significant predicted structure over the PUM1/2 binding site. Among these variants, variants that contained significant predicted structure outside the PUM1/2 site were defined as those having predicted ensemble folding free energy below $-0.5$ kcal/mol, with PUM1/2 site constrained to single-stranded state in RNAfold. Predicted structures for three variants with the greatest observed weakening of binding are shown, with the PUM1/2 consensus site highlighted in yellow; the structures were generated using the RNAfold web server and represent the MFE structures. While the majority of effects were less than 1 kcal/mol, the data suggest that stable base-pairing in the vicinity of the PUM1/2 site may slightly weaken binding, likely due to steric clashes with the protein.

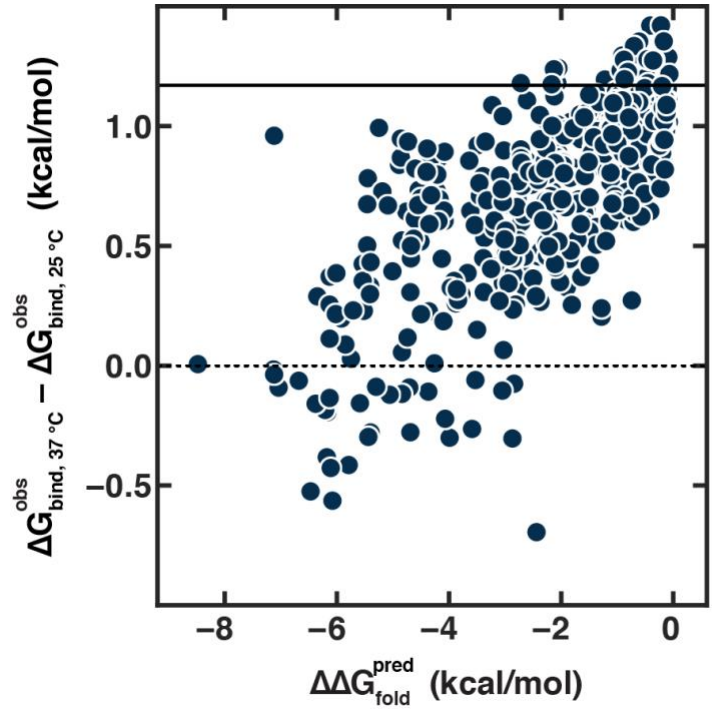

**Fig. S5.** Comparison of the difference between the measured affinity ($\Delta G$) of consensus (UGUAUAUA) containing RNAs at 37 °C and 25 °C and the structural stability predicted by RNAfold. The solid black line indicates the difference between the measured affinities for unstructured RNAs ($\Delta G_{\text{fold}}^{\text{pred}} > -0.2$ kcal/mol) containing consensus (UGUAUAUA) binding sites at 37 °C and 25 °C. The difference between the observed binding affinities at 37 °C and 25 °C decreases as the predicted structure becomes more stable because RNA secondary structure is destabilized at higher temperatures. Some RNA variants with the most stable predicted secondary structure actually bind PUM2 with higher affinity at 37 °C rather than 25 °C (y-axis < 0) because the weakening of PUM2 binding at higher temperature is completely offset by weakening of RNA stability.

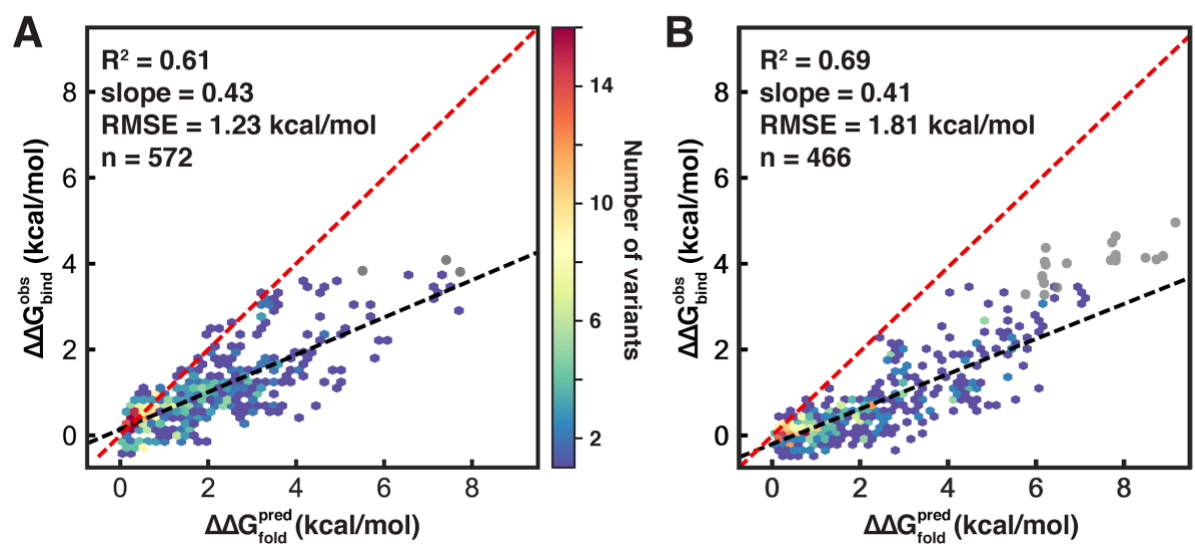

**Fig. S6.** (A, B) Scatterplot of predicted and observed structure effects on PUM2 binding to mutant (UGUAUAUU) sites (A) and on PUM1 binding to consensus (UGUAUAUA) sites (B). The predicted line is shown in red, and the best fit line is shown in black. Grey points indicate variants with affinities below the threshold for high-confidence affinity determination.

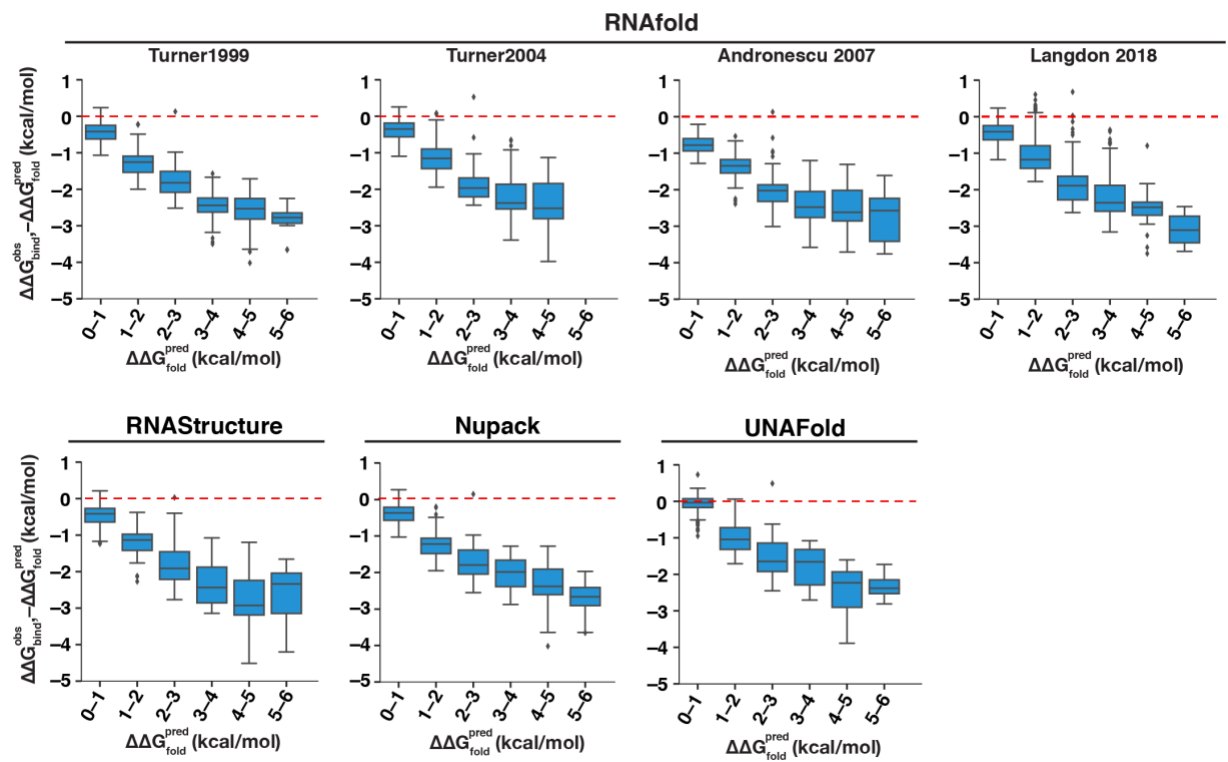

**Fig. S7.** Differences between predicted and observed structure effects across bins of predicted structure effects. The comparisons are shown for the different NN prediction packages and parameter sets. The plotted values correspond to measured and predicted structure effects at 37 °C. The red line indicates perfect agreement with predicted stabilities (corresponding to the diagonal in Figure 3A). Only bins with at least 10 variants are shown.

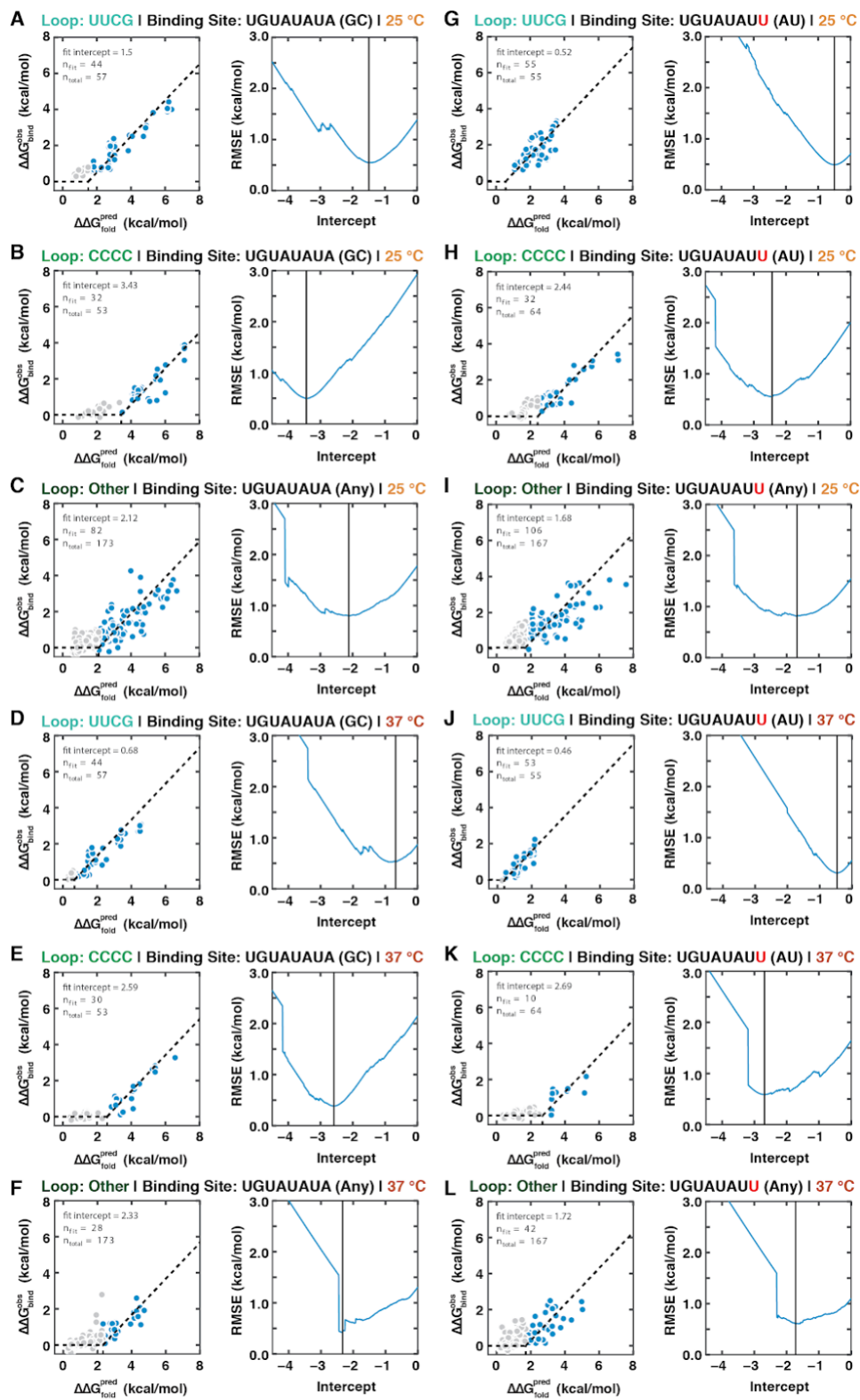

**Fig. S8.** Results of fitting lines with a slope of 1 to determine loop stability offsets. The displacement (horizontal dashed line) was determined by finding the intercept at which fitting a line with slope 1 to all points above the intercept minimized the least squared error. The best fit

intercepts to the predicted and observed $\Delta\Delta G$ are shown at the left of each panel and the relationship between the stability offset and the RMSE is shown at the right of each panel. The points for which the predicted RNA stability was greater than the intercept that produced the lowest RMSE are blue and the other points are gray. The panels are divided by the PUM binding site sequences (consensus UGUAUAUA vs. mutant UGUAUAU<u>U</u>), the temperature of the experiment (25 °C vs 37 °C—orange vs. red) and the loop at the end of the hairpin (UUCG vs. CCCC vs. all other loops—shades of green).

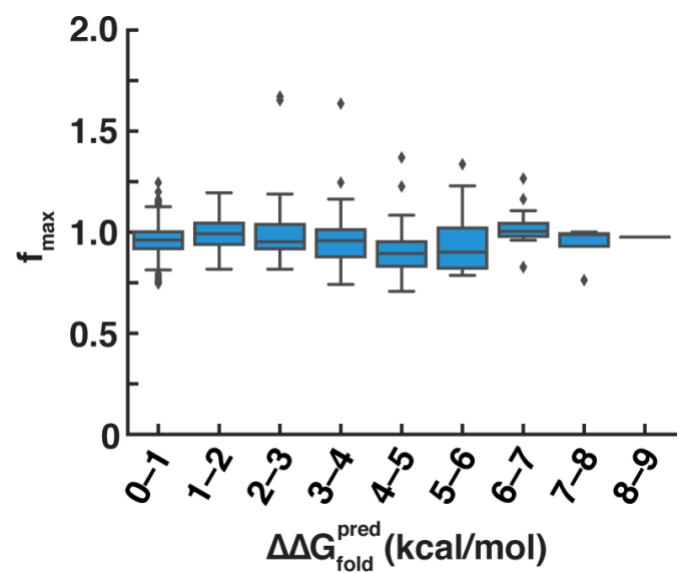

**Fig. S9.** Distribution of fit $f_{\max}$ values across predicted structure stability bins.

### Supplemental Tables

**Table S1:** Numbers of publications referencing major structure prediction algorithms.

| Algorithm | Publication | Number of citations* |
| --- | --- | --- |
| mfold/UNAFold | (15) | 10731 |
|  | (12) | 824 |
| Vienna RNA | (16) | 2422 |
|  | (17) | 2044 |
|  | (18) | 1139 |
|  | (1) | 1497 |
| RNAstructure | (7) | 1241 |
|  | (19) | 292 |
|  | (20) | 822 |
|  | (21) | 114 |
| NUPACK | (11) | 574 |
| Sfold | (22) | 458 |
| RNAsoft | (23) | 137 |

\* Number of Google Scholar citations on 12/14/18

**Table S3.** Offsets from predicted values for different hairpin loops and closing base pairs across prediction algorithms (see Figure 5).

|  |  | Fit offset from predicted values (kcal/mol) |  |  |  |  |  |
| --- | --- | --- | --- | --- | --- | --- | --- |
| Loop |  | UUCG |  | CCCC |  | Other |  |
| Closing bp |  | AU | GC | AU | GC | AU | GC |
| n |  | 69 | 49 | 82 | 50 | 113 | 5 |
| Program | RNAstructure | 1.23 | 2.26 | 2.39 | 1.31 | 3.03 | 3.38 |
|  | UNAFold | 0.48 | 2.40 | 1.80 | 1.05 | 2.20 | 2.68 |
|  | Nupack | 1.01 | 2.75 | 2.09 | 1.40 | 2.36 | 3.16 |
|  | RNAfold | 1999 | 1.04 | 2.76 | 2.68 | 2.74 | 2.48 |
|  |  | 2004 | 0.64 | 0.99 | 2.68 | 2.64 | 2.54 |
|  |  | 2007 | 0.92 | 1.98 | 2.25 | 2.73 | 2.40 |
|  |  | 2018 | 0.59 | 0.33 | 2.73 | 2.51 | 2.47 |

**Table S4:** Numbers of library variants passing each of the filters. The numbers indicate how many variants (and what percentage of originally designed variants) met the indicated criterion and all of the preceding criteria in the table. Numbers are shown for PUM2 binding at 25 °C.

|  | UGUAUAUA | % | UGUAUAUU | % |
| --- | --- | --- | --- | --- |
| **Designed** | 1297 |  | 1362 |  |
| **Measured (≥5 clusters)** | 1189 | 91.7 | 1262 | 92.7 |
| **95% CI < 1 kcal/mol** | 1177 | 90.7 | 1252 | 91.9 |
| **ΔG < −8.5 kcal/mol** | 1144 | 88.2 | 1235 | 90.7 |
| **No alternative sites\*** | 686 | 52.9 | 692 | 50.8 |
| **No structure in bound state\*\*** | 486 | 37.5 | 572 | 42.0 |

\* No alternative non-overlapping sites with predicted $\Delta\Delta G < 5$ kcal/mol and no more than one UGUA site.

\*\* $\Delta G_{\text{fold}}^{\text{pred}} > -0.5$ kcal/mol after constraining the PUM2 site to single-stranded state.
